# Inhibition of USP28 overcomes Cisplatin-Resistance of Squamous Tumors by Suppression of the Fanconi Anemia Pathway

**DOI:** 10.1101/2020.09.10.291278

**Authors:** Cristian Prieto-Garcia, Oliver Hartmann, Michaela Reissland, Thomas Fischer, Carina R. Maier, Mathias Rosenfeldt, Christina Schülein-Völk, Kevin Klann, Reinhard Kalb, Ivan Dikic, Christian Münch, Markus E. Diefenbacher

## Abstract

Squamous cell carcinomas (SCC) frequently have a limited response to or develop resistance to platinum-based chemotherapy, and have an exceptionally high tumor mutational burden. As a consequence, overall survival is limited and novel therapeutic strategies are urgently required, especially in light of a rising incidences. SCC tumors express ΔNp63, a potent regulator of the Fanconi Anemia (FA) DNA-damage response pathway during chemotherapy, thereby directly contributing to chemotherapy-resistance. Here we report that the deubiquitylase USP28 affects the FA DNA repair pathway during cisplatin treatment in SCC, thereby influencing therapy outcome. In an ATR-dependent fashion, USP28 is phosphorylated and activated to positively regulate the DNA damage response. Inhibition of USP28 reduces recombinational repair via an ΔNp63-Fanconi Anemia pathway axis, and weakens the ability of tumor cells to accurately repair DNA. Our study presents a novel mechanism by which tumor cells, and in particular ΔNp63 expressing SCC, can be targeted to overcome chemotherapy resistance.

**Significance:** Limited treatment options and low response rates to chemotherapy are particularly common in patients with squamous cancer. The SCC specific transcription factor ΔNp63 enhances the expression of Fanconi Anemia genes, thereby contributing to recombinational DNA repair and Cisplatin resistance. Targeting the USP28-ΔNp63 axis in SCC tones down this DNA damage response pathways, thereby sensitizing SCC cells to cisplatin treatment.

## INTRODUCTION

Squamous tumors arise in various tissues, including skin, esophagus, pancreas, cervix, head, neck and lung, and are among the most mutated cancer entities, as identified by Next Generation Sequencing(Bray et al., 2018; Cancer Genome Atlas, 2015; Cancer Genome Atlas Research, 2012). Despite novel and detailed insights into the genetics driving this cancer subtype, treatment options are rather limited and predominantly restricted to conventional (DNA damage inducing chemotherapy) or personalized approaches, such as receptor tyrosine kinases (RTK) and immune checkpoint inhibitors(Drilon et al., 2018; Ettinger et al., 2017; Kim et al., 2018; Mok et al., 2009; Palyca et al., 2014; Parashar et al., 2013; Stratigos et al., 2015).

Although initial treatment responses in patients are observed, tumors frequently develop strategies to overcome therapy-induced challenges(Saleh et al., 2019a; Saleh et al., 2019b; Stewart and Abrams, 2008). Tumor cells achieve this by either acquiring additional mutations within the initially targeted tumor-essential pathway(s) or by activating alternative signaling cascades restoring proliferation advantages(Boussemart et al., 2016; Khaliq and Fallahi-Sichani, 2019; Noeparast et al., 2019; Zaman et al., 2019). Examples of this cancer adaptation were observed in patients treated with RTK inhibitors specifically targeting the hotspot mutation L858R within the *EGFR-gene* (*EGFR^L858R^*)(Liu et al., 2018). Despite early treatment responses, tumors quickly recur and Next Generation Sequencing identified the presence of a novel mutation within *EGFR, T790M*, thereby negating the inhibitor and rendering the receptor constitutively active(Nukaga et al., 2017; Zhou et al., 2019).

To prevent the ability of cancer cells to escape treatment via alternative pathways, one attractive therapeutic strategy is the direct targeting of important downstream effector molecules and essential proto-oncogenes, as tumor cells, in contrast to non-transformed cells, are more susceptible to changes in the abundance or activity of these proteins(Chen et al., 2018; Orlando et al., 2019). Due to the oncogene driver heterogeneity observed in SCC, targeting a commonly expressed nominator found within SCC is interesting, as this means that a widely applicable therapeutic strategy for SCC could be developed. One such factor, distinguishing SCC from other tumor entities, is the proto-oncogene ΔNp63. In contrast to adenocarcinomas, SCC tumors express and are dependent on ΔNp63, as SCC require it to maintain an epidermal lineage identity(Ratovitski, 2014; Romano et al., 2012). Ectopic expression of ΔNp63 is able to drive ‘trans-differentiation’ and impose an epidermal signature, thus demonstrating the potency of this transcription factor as the master regulator of SCC formation(Hamdan and Johnsen, 2018; Soares and Zhou, 2018; Somerville et al., 2018), and that the genetic depletion of ΔNp63 is detrimental to SCC *in vivo(Ramsey et al., 2013; Romano et al., 2012*).

Not only do SCC require ΔNp63 as a marker protein and master regulator of SCC lineage and identity, but it also contributes to the chemotherapy resistance phenotype observed in SCC. Modulation of ΔNp63 protein abundance was sufficient to resensitize cells to platin-based therapy(Matin et al., 2013) and can be attributed to its ability to regulate the expression of DNA-damage response (DDR) genes(Lin et al., 2009). The Fanconi Anemia pathway(Bretz et al., 2016) is the foremost DDR pathway directly regulated by ΔNp63 that modulates platin-based treatment responses in SCC. ΔNp63 is recruited to several key genes of this pathway and drives their expression, specifically during DDR-inducing therapy(Bretz et al., 2016).

A weakness of SCC is its dependence for the expression of the deubiquitylase USP28(Prieto-Garcia et al., 2020a). USP28 regulates ΔNp63 protein abundance during SCC tumor initiation and is required for tumor maintenance. Targeting USP28 in SCC is a valid strategy, as the first-generation small molecule inhibitor AZ1’s inhibition of USP28 suppressed tumor growth in a murine isogenic transplant model (Prieto-Garcia et al., 2020a). USP28 regulates the abundance of proto-oncogenes and promotse proliferation of cancer cells. Furthermore, it is also involved in chromatin stability, segregation and DNA damage signaling and response(Diefenbacher et al., 2015; Diefenbacher et al., 2014; Fong et al., 2016; Lambrus et al., 2016; Li et al., 2019; Meitinger et al., 2016; Muller et al., 2020; Popov et al., 2007b; Schulein-Volk et al., 2014; Wang et al., 2018; Zhang et al., 2015; Zhao et al., 2018). Its role in the DDR pathway, however, is unclear(Knobel et al., 2014).

Here we report that the deubiquitylase USP28 regulates, via ΔNp63, the maintenance of chromatin/DNA integrity during DNA damaging chemotherapy with Cisplatin. USP28 is activated in an ATR dependent fashion, stabilizes itself, MYC and ΔNp63, respectively. Loss of USP28 induces a pro-DNA damage signature, while weakens the ability of tumor cells to maintain a functional DNA damage program and accurately repair DNA. This mechanism provides a novel target to sensitize in particular ΔNp63 positive SCC by chemotherapy.

## RESULTS

### USP28 is Recruited to Sites of DNA Damage and Phosphorylated by ATR, Not ATM, Upon Cisplatin Treatment

Previous studies demonstrated that USP28 is recruited to DNA damage upon exposure to ionizing radiation(Knobel et al., 2014; Popov et al., 2007a; Zhang et al., 2006). As chemotherapeutic agents, such as Cisplatin, are the mainstay compound for chemotherapy of SCC, we considered whether USP28 is recruited to DNA damage loci induced by DNA crosslinking agents. The human SCC line A431 was exposed to either DMSO or 5μM CPPD (Cisplatin) for 6 hours, followed by immunofluorescence staining against USP28 and DNA damage markers (Figure 1A). While USP28 was evenly distributed throughout the nucleus in control cells, USP28 formed nuclear foci similar to the DNA damage markers ɣ-H2AX, TP53BP1, p-ATM or p-ATR upon CPPD exposure (Figure 1A). Next, we performed CPPD pulse chase experiments to address if the amount of chromatin-associated USP28 is altered in CPPD-treated cells. A431 cells were treated with either DMSO or CPPD (5μM) and samples collected at indicated time points (Figure 1B). In non-stimulated cells (time point 0), low amounts of USP28 was bound to chromatin, which increased during CPPD treatment in a time-dependent fashion (Figure 1B). We then investigated if USP28 is modified upon CPPD treatment by ATM or ATR, as it contains a conserved ATM/ATR substrate SQ/TQ motive (Figure 1C and S1A). A431 cells were exposed to CPPD for 24 hours. USP28 phosphorylation upon CPPD treatment was confirmed using antibodies recognising serine 67 and serine 714 phosphorylation on USP28 (Figure 1D). Increasing concentrations of CPPD resulted in an increase in overall USP28 phosphorylation (Figure 1D). Exposure to CPPD also resulted in an increase in USP28 activity, as measured by ubiquitin-suicide probe assay (Figure 1E). The increased deubiquitylase-activity of USP28 led to an enhanced deubiquitylation of USP28 substrates, such as c-MYC and ΔNp63, as measured by <u>t</u>andem <u>u</u>biquitin <u>b</u>inding <u>e</u>ntity (TUBE) pull down (Figure 1F). It is important to note that the USP28 substrates (ΔNp63, c-MYC, c-JUN) were upregulated in control A-431 but not in USP28-depleted cells upon CPPD treatment (Figure S1B), demonstrating that USP28 is indeed required to stabilise these factors during DNA damage induction. Furthermore, USP28 mutant lung cancer cell lines H23 (adenocarcinoma, ADC) and Sk-Mes1 (SCC) (Figure S1C, S1D and S1E) failed to stabilize c-JUN or c-MYC protein abundance upon CPPD exposure (Figure S1C, S1D and S1E). Activation of the DNA damage response was observed, indicated by ɣ-H2AX phosphorylation (Figure S1E).

**Figure 1.**
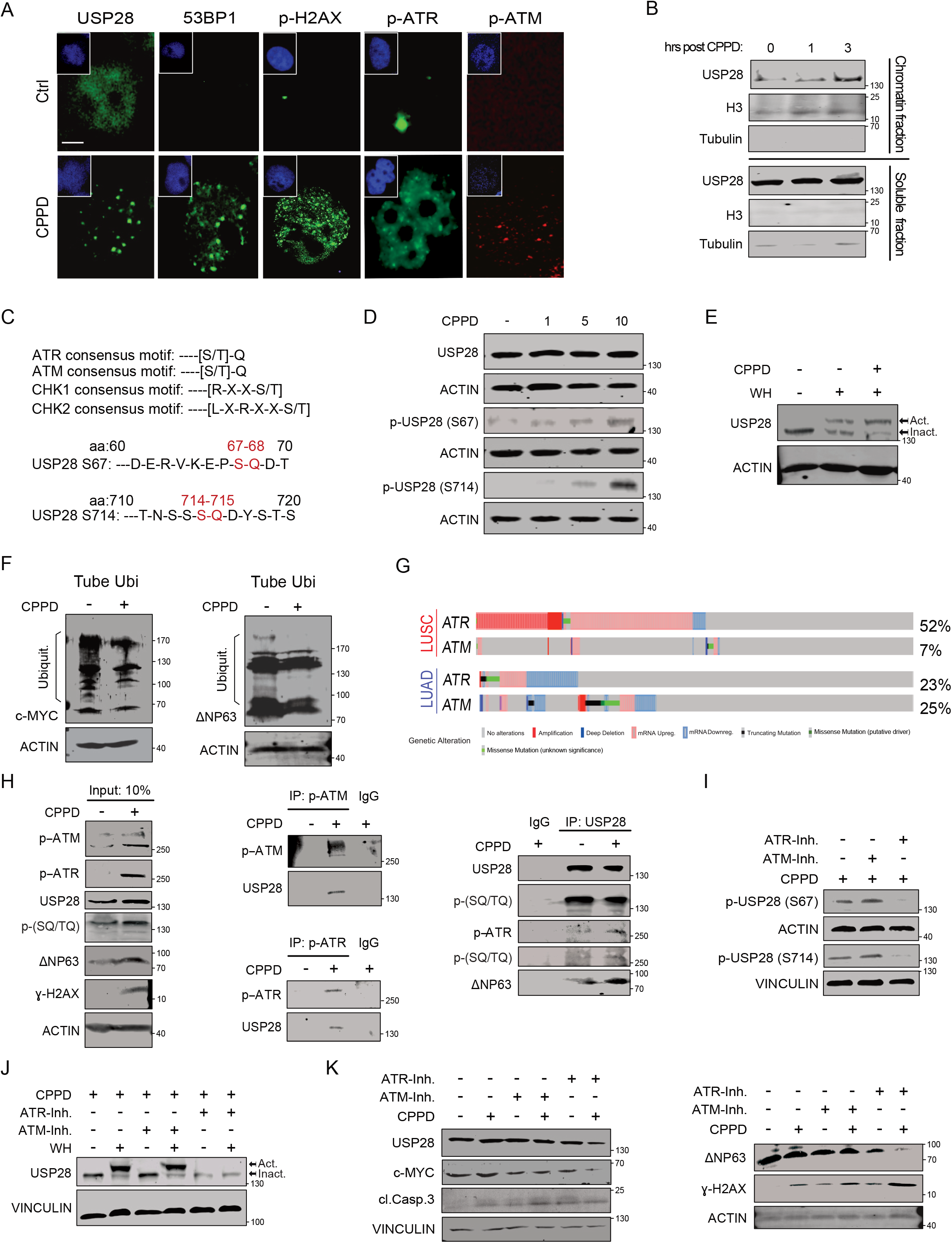
USP28 is recruited to DNA damage sites and phosphorylated by ATR, not ATM, upon Cisplatin treatment. A) Immunofluorescence staining of endogenous USP28, 53BP1, ɣ-H2AX, phospho-ATR and phospho-ATM in A431 cells exposed to either DMF or 5μM Cisplatin for 6 hours. DAPI as nuclear counterstain. Scale bar= 10μm. n=3. B) Chromatin and nucleoplasm fractionation, followed by immunoblotting of endogenous USP28 in A431 cells exposed to 5μM CPPD at indicated time points. Histone H3 and Tubulin serve as loading control. n=3. C) Immunoblotting of total and phosphorylated USP28 at serine 67 and 714 in A431 cells exposed to indicated concentrations of CPPD for 6 hours. n=3. D) Ubiquitin suicide probe (warhead) assay, followed by immunoblotting against USP28 in A431 cells exposed to 5 μM CPPD for 6 hours. ‘Act.’ arrow indicates active USP28.’ Inact.’ arrow indicates inactive USP28. ACTIN serves as loading control. n=3. E) Tandem-ubiquitin binding entity (TUBE) pulldown of endogenous ubiquitin, followed by immunoblotting against endogenous c-MYC and ΔNp63 in control of 5μM CPPD treated A431 cells for 6 hours. ACTIN serves as loading control. n=3. F) ATM/ATR SQ/TQ consensus motive alignment of human CHK1, CHK2 and USP28 G) Genetic alteration of ATR and ATM in lung SCC (LUSC) and lung ADC (LUAD) tumors. Publicly available patient data obtained from CBIOPORTAL (www.cbioportal.org) H) Immunoprecipitation of control rabbit IgG, endogenous phospho-ATM, phospho-ATR or USP28 of either control or 5μM CDD treated A431 cellsfor 6 hours, followed by immune-blotting against ATM, ATR, USP28, phospho-H2AX, ΔNp63 or SQ/TQ motive specific antibodies. ACTIN serves as loading control. n=3. I) Immunoblotting of phosphorylated USP28 at serine 67 and 714 in A431 cells exposed to 5 μM CPPD for 6 hours and co-treatment with 15 μM ATM kinase inhibitor KU55933 or 2.5 μM ATR kinase inhibitor VE 821. n=3. J) Ubiquitin suicide probe (warhead) assay, followed by immunoblotting against USP28 in A431 cells exposed to 5 μM CPPD for 6 hours and co-treatment with 15 μM ATM kinase inhibitor KU55933 or 2.5 μM ATR kinase inhibitor VE 821. Act. arrow indicates active USP28. Inact. arrow indicates inactive USP28. ACTIN serves as loading control. n=3. K) Immunoblotting against endogenous USP28, c-MYC, Cleaved Caspase 3, ΔNp63 and phospho-H2AX in A431 cells treated with either DMF or 5μM CPPD for 24 hours and co-treatment with 15 μM ATM kinase inhibitor KU55933 or 2.5 μM ATR kinase inhibitor VE 821. ACTIN and VINCULIN serve as loading control. n=3. See also Figure S1

We next asked whether ATM or ATR phosphorylate and activate USP28 upon CPPD treatment. Based on publically available datasets, ATR is frequently upregulated or amplified in SCC when compared to ADC, while ATM is commonly downregulated or lost (Figure 1G and S1F). Immunoprecipitation of endogenous phosphorylated ATM and ATR was able to co-immunoprecipitate endogenous USP28 in A431 cells treated with 5uM CPPD for 6 hours, while no interaction was detected in untreated cells (Figure 1H). Conversely, endogenous USP28 co-immunoprecipitated with phosphorylated ATR, along with ɣ-H2AX and ΔNp63, in cells exposed to CPPD (Figure 1H). Notably, USP28 interacts with ΔNp63 stronger after CPPD treatment (Figure 1H). Next, we treated A431 cells with CPPD and co-treated with either the ATM inhibitor KU55933 or the ATR inhibitor VE-821 for 24 hours. While treatment with CPPD-induced USP28 phosphorylation, inhibition of ATM did not alter the protein abundance or the phosphorylation of USP28 (Figure 1I). Co-treatment with VE-821 in CPPD-treated cells abolished the phosphorylation of USP28 at its conserved SQ/TQ sites (Figure 1I). Deubiquitylase activity assays in the presence of CPPD and ATM or ATR inhibitors revealed that ATM inhibition did not alter the total amount nor active state of USP28 (Figure 1J), while ATR inhibition resulted in an inhibition of USP28 and led to the reduction of overall USP28 abundance (Figure 1J). A431 cells exposed to CPPD upregulate the protein abundance of c-MYC and ΔNp63, alongside increased abundance of ɣ-H2AX (Figure 1K). This increase is required to resolve DDR stress. Exposure to the ATM inhibitor showed reduced protein abundance of c-MYC and an increase in ɣ-H2AX, which was further increased during CPPD treatment (Figure 1K). Cells undergo DDR stress, as cleaved caspase 3 was enriched in KU55933/CPPD cotreated cells. Exposure to VE-821showed reduced amounts of USP28 and its substrates c-MYC and ΔNp63. Upon co-exposure to CPPD, USP28 protein abundance was further reduced, together with the oncoproteins c-MYC and ΔNp63 (Figure 1K). VE-821/CPPD co-treatment resulted in an increased cleavage of caspase 3 and increased ɣ-H2AX abundance (Figure 1K).

These data indicate that USP28 is recruited to DNA damage sites upon exposure to cisplatin and its interaction with ATR. ATR-mediated phosphorylation enhances USP28 activity, which facilitates the stabilization of pro-survival factors, such as c-MYC and ΔNp63, to counteract DDR stress induced by CPPD. USP28, therefore, is an important player within the ATM-/ATR-pathway and operates downstream of ATM (during IR induced double strand breaks) or ATR.

### Phosphorylation of USP28 upon Cisplatin exposure is required to repair DNA damage in SCC

USP28 is recruited to DNA damage sites and is phosphorylated by ATM (IR-dependent) or ATR (platin-induced) at two conserved SQ/TQ motive sites, serine 67 and serine 714 (Figure S1A and 2A). The role of USP28 phosphorylation is poorly understood, but has been associated with DNA repair and apoptosis(Zhang et al., 2006). To examine the role of phosphorylated USP28 in A431 cell lines we generated phospho-mutant knock-in cell lines for serine 67 (S67A), serine 714 (S714A), or both simultaneously (S67/714A) by using CRISPR/Cas9 (Figure 2B and 2C). A similar approach was conducted using the human embryonic, non-tumor, kidney cell line HEK293 (Figure S2A). CRISPR-targeted cell clones were propagated and subjected to CPPD treatment for 6 hours, followed by western blotting against USP28 and the SQ/TQ epitope antibody (Figure 2B and S2A). While total protein amounts of USP28 were not affected in A431 cells, endogenous targeting of serine 67 or 714, or both, diminished the phosphorylation at the SQ/TQ sites within USP28 (Figure 2B and S2A). This was further confirmed by using the phosphosite specific antibodies of USP28 (Figure 2C and S2A). It is worth noting that targeting of one phosphosite, either serine 67 or serine 714, diminished the phosphorylation of the second site. This suggests that either site might be required to facilitate the second phosphorylation event. In line with previous experiments, endogenous mutation of the SQ/TQ motives reduced overall USP28 deubiquitylase activity upon CPPD exposure, as measured by ubiquitin suicide probe assays (Figure 2D and S2B).

**Figure 2.**
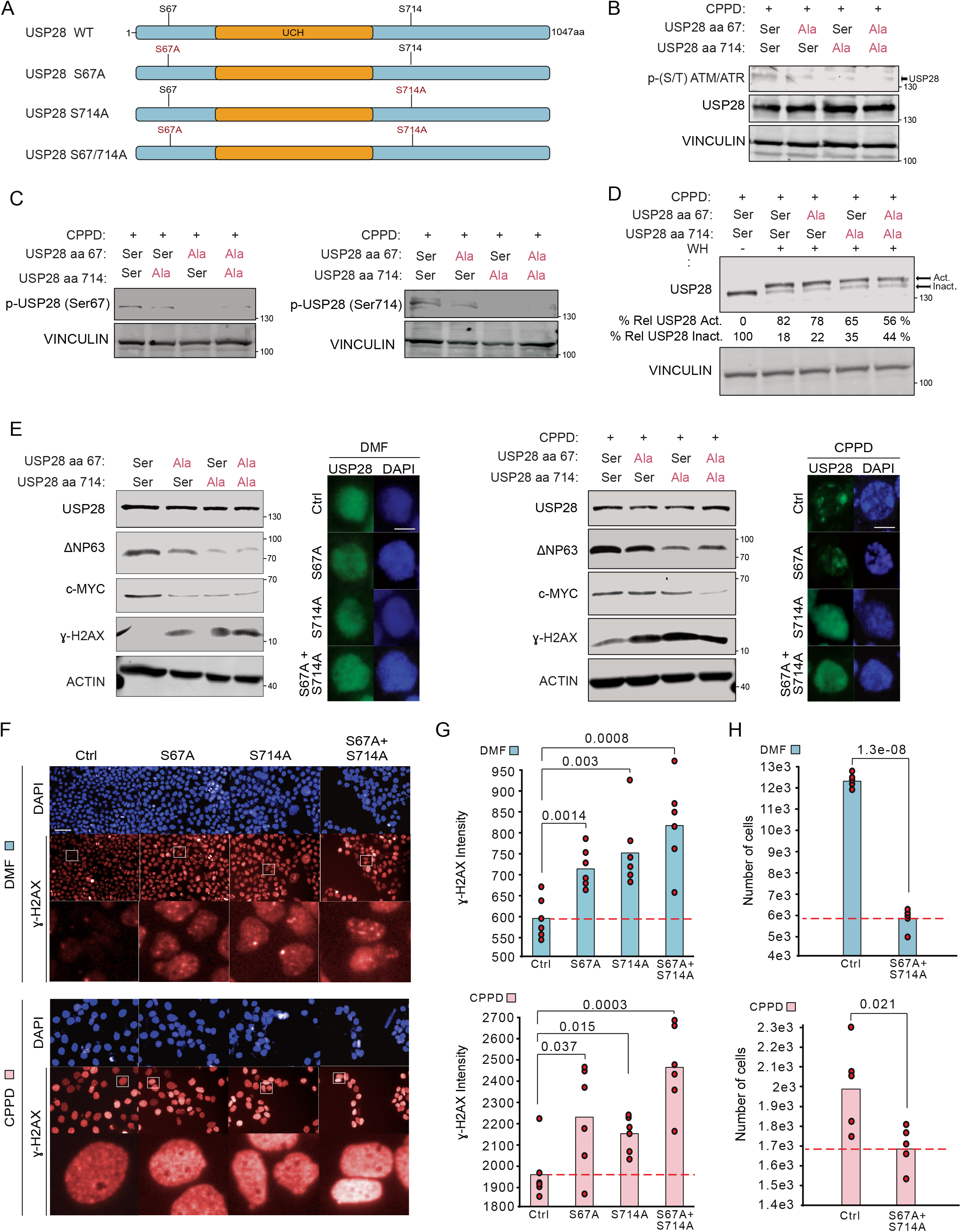
Phosphorylation of USP28 upon Cisplatin exposure is required to repair DNA damage in SCC. A) Schematic representation of the point mutations introduced into USP28 in A431 and HEK293T cell lines. Red=mutation; Black=WT B) Immunoblotting against endogenous USP28 and ATR/ATM SQ/TQ motif in control, S67A, S714A and S67A+S714A mutant A431 cells treated with 5μM CPPD for 6 hours. VINCULIN serves as loading control. n=3 C) Immunoblotting of phosphorylated USP28 at serine 67 and 714 in control, S67A, S714A and S67A+S714A mutant A431 cells treated with 5μM CPPD for 6 hours. VINCULIN serves as loading control. n=3 D) Ubiquitin suicide probe assay, followed by immunoblotting against USP28 in control, S67A, S714A and S67A+S714A mutant A431 cells exposed to 5 μM CPPD for 6 hours. ‘Act.’ arrow indicates active USP28.’ Inact.’ arrow indicates inactive USP28. VINCULIN serves as loading control. n=3. E) Immunoblotting of USP28, c-MYC, ΔNp63 and ɣ-H2AX in control, S67A, S714A and S67A+S714A mutant A431 cells. ACTIN serves as loading control. n=3. Immunofluorescence against endogenous USP28 in control, S67A, S714A and S67A+S714A mutant A431 cells. Scale bar= 10μm DAPI served as nuclear marker. n=3. Cells were either treated with DMF or 5 μM CPPD for 6 hours. F) Immunofluorescence against endogenous phosho-H2AX in control, S67A, S714A and S67A+S714A mutant A431 cells, treated with either DMF (blue) or 5μM CPPD (pink) for 48 hours. DAPI served as nuclear marker. n=6. Phospho-H2AX intensity was calculated measuring 15 fields per well (n=6). Scale bar= 1 00μm. P-values were calculated using two-tailed T-test statistical analysis. G) Number of cells in control and S67A+S714A mutant A431 cells, treated with either DMF (blue) or 5μM CPPD (pink) for 48 hours. Number of cells were calculated measuring 15 fields per well (n=5). P-values were calculated using two-tailed T-test statistical analysis. See also Figure S2

We next assessed downstream effects of blocking USP28 phosphorylation in the knock-in cell lines by western blotting against the USP28 substrates c-MYC and ΔNP63 (Figure 2E). In non-treated wild type cells c-MYC and ΔNP63 were readily detectable, upon mutation of serine 67 to alanine; however, total protein abundance of c-MYC and ΔNP63 were significantly reduced, along with an increase in phosphorylation of TP53 (serine 15) and ɣ-H2AX (Figure 2E and S2C). This was further increased in cells harboring mutations at serine 714, or both (Figure 2E and S2C).

When cells were exposed to CPPD, mutant knock in cell lines failed to stabilize c-MYC and ΔNP63, but increased phosphorylation of ɣ-H2AX (Figure 2E). Mutation of USP28 at serine 67 (S67A) only showed a mild reduction in c-MYC and ΔNP63 when compared to S714A or S67/714A mutant cells. Next, we performed immunofluorescence staining against endogenous USP28 and compared control, S67A, S714A and S67/714A clones in control or 6 hours CPPD treatment (Figure 2E). In solvent-treated A431 cells, USP28 was evenly distributed throughout the nucleus (Figure 2E). Upon treatment with CPPD, control as well as S67A-mutant A431 clones showed USP28 foci within the nucleus and DNA compaction, as seen by DAPI (Figure 2E), indicating that loss of serine 67 phosphorylation does not alter the ability of USP28 recruitment to DNA damage sites. Mutation of serine 714 to alanine abolished the presence of USP28 foci upon CPPD exposure. Similar results were obtained when analysing the double knock-in mutants (Figure 2E), hinting to a dominant role of serine 714.

Mutation of the SQ/TQ motives within USP28 induced basal replication and DDR stress, as seen by persistent ɣ-H2AX foci formation under physiological conditions, when compared to parental control cells (Figure 2F and 2G). Upon CPPD treatment, mutant cells showed a significant increase in ɣ-H2AX foci formation (Figure 2F and 2G). Similar effects were seen with the DNA damage marker TP53BP1 in mutant A431 cells under basal and CPPD treated conditions (Figure S2D). Cell proliferation was significantly reduced in cells carrying mutations at S714A or S67/714A, which ultimately resulted in clonal loss in a long term culture (Figure 2H and S2E).

These findings strongly suggest that impairing USP28 phosphorylation increases the occurrence of DDR lesions and reduces cell proliferation and/or viability.

### Loss of USP28 Negatively Affects the Expression of DDR Effector Proteins in SCC

Previous studies have shown that SCC tumors exhibited limited response rates to therapy and consequently poorer prognosis than ADC in overall survival(Prieto-Garcia et al., 2020a; Ruiz et al., 2019). In lung SCC, the expression of DDR signature genes was elevated when compared to normal tissue or ADC (Figure S3A). Analysis of the top expressed genes correlating with poor prognosis (red marks) revealed that several DDR genes, such as FANCI, PCNA or RFC4, were upregulated in SCC when compared to ADC (Fig S3B). Furthermore, enhanced expression of DDR signature genes coincided with shortened overall survival (Figure 3A). Specifically, under chemotherapy, the relative expression of DDR signature genes is an indicative prognostic marker, as elevated expression strongly correlates with poor survival, based on publicly available datasets (Figure 3A, right panel).

**Figure 3.**
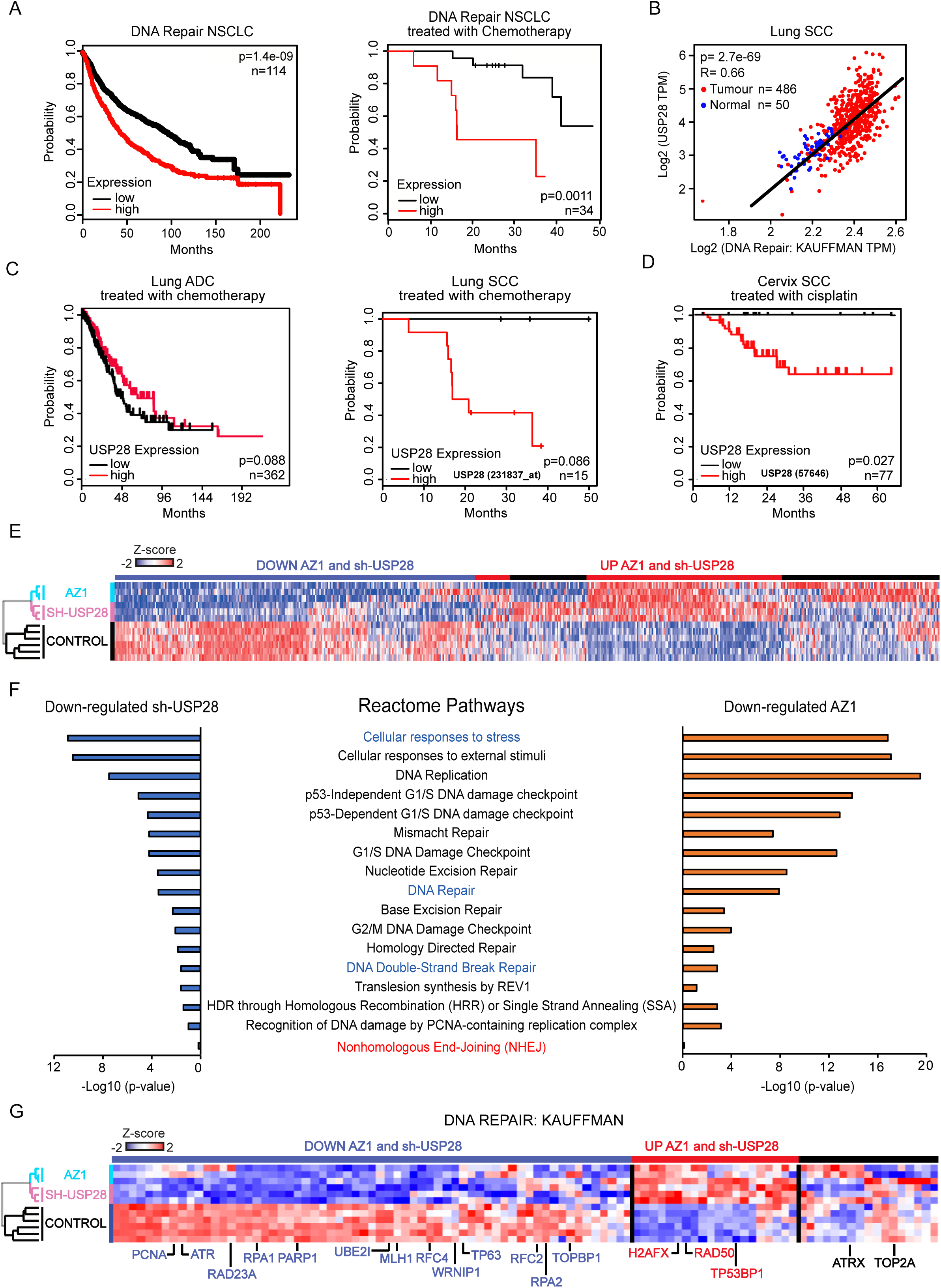
Loss of USP28 negatively affects the expression of DDR effector proteins in SCC. A) Public available patient survival data of NSCLC patients are stratified by relative expression of DNA damage gene expression according to Kauffmann signature selection. Left panel= All NSCLC patients; Right panel= Only NSCLC treated with chemotherapy. Generated with the open source tool www.kmplot.com. B) Correlation between USP28 and Kauffmann DNA repair signature gene expression in human lung SCC and normal tissue. Spearman correlation R=0.66, p=2.7e-69. Generated with the open source tool www.gepia2.cancer-pku.cn. C) Public available patient survival data of lung ADC and SCC cancer patients treated with chemotherapy and stratified by relative expression of USP28. Left panel= ADC patients; Right panel= SCC patients. Generated with the online tools www.kmplot.com and www.r2.amc.nl. D) Publicly available patient survival data of cervix SCC cancer patients treated with cisplatin (CPPD) and stratified by relative expression of USP28. Generated with the online tools www.r2.amc.nl. E) Heatmap of whole cell proteome analysis of A431 cells treated with the DUB inhibitor AZ-1 or DMSO (Control), shRNA targeting USP28 or Non-targeting control NTC (Control). n=3. F) Reactome pathway analysis of proteomic data upon genetic depletion or pharmacological targeting by AZ-1 of USP28 in A431 cells. Highlighted are pathways involved in DNA damage signalling, response and clearance. Generated with the open source tool www.pantherdb.org. NHEJ (red) was not affected upon AZ-1 or USP28 genetic depletion. G) Heatmap of proteome analysis according to the Kauffmann DNA damage signature of A431 cells treated with the DUB inhibitor AZ-1 or DMSO control), shRNA targeting USP28 or Non-targeting control NTC. Blue= Down-regulated in AZ1/sh-USP28; Red=Up-regulated in AZ1/sh-USP28. n=3. See also Figure S3

As USP28 is recruited to sites of DNA damage and is activated by ATR and ATM, we considered if USP28 is involved in chemotherapy resistance. By analyzing publicly available datasets, we could identify a strong correlation between DDR gene signatures and USP28 expression, particularly in lung SCC, when compared to non-transformed lung samples (Figure 3B). Stratifying patient survival datasets for lung ADC and SCC with regard to chemotherapy and USP28 expression highlighted a strong correlation with significantly shortened patient survival in SCC (Figure 3C). Similar observations were obtained when we analyzed a publicly available dataset of cervix SCC survival data upon cisplatin treatment (Figure 3D). Patients with USP28^high^ expressing tumors had a significantly shortened overall survival rate, when compared to a USP28^low^ cohort.

To address the potential involvement of USP28 in chemoresistance, we assessed the impact of USP28 depletion on DDR protein abundance. USP28 was thus either silenced in A431 cells by an inducible shRNA sequence, or the cells were treated with AZ1, a dual specific USP25/28 inhibitor(Wrigley et al., 2017). Changes in protein abundance were measured by whole proteome mass spectrometry (Figure S3C and S3D). Silencing or inhibition of USP28 by AZ1 resulted in comparable changes of the whole proteome when compared to control (non-targeting shRNA or DMSO, Figure 3E, S3C and S3D). Loss of USP28 activity predominantly affected cellular pathways associated with stress, cell cycle progression and DNA damage checkpoint and repair (Figure 3F and S3E). Furthermore, proteins involved in DNA replication were significantly reduced upon interference with USP28 (Figure 3F). Of note, USP28 impairment and downregulation of its substrate ΔNp63 caused upregulation of certain DNA repair associated proteins, such as TP53BP1 or RAD50 and downreagulation of proteins involved in DNA recombinational repair, others, such as ATR, PCNA or WRNIP1, and specially proteins related to DNA, such as RAD51, ATR, RPA1, RPA2 PCNA or WRNIP1 (Figure 3G and S3F).

Loss of USP28, by genetic depletion or inhibition of the catalytic activity, deregulated the replication machinery and downregulated the DDR signalling in the SCC cell line A431.

### USP28-ΔNp63 Axis is Required for DDR Upon Cisplatin Treatment and Chemoresistance in SCC

Since USP28^high^ tumors are associated with chemo-resistance we wondered if inactivation or lack of USP28 could sensitize SCC cells to chemotherapeutic drugs. Another protein, ΔNp63, has been shown to influence the expression of DDR genes, specifically in SCC, making an USP28-ΔNp63 axis a reasonable target(Bretz et al., 2016; Prieto-Garcia et al., 2020b). To assess if USP28 is a limiting factor in chemoresistance, we over-expressed USP28 in a chemosensitive cell line, BEAS-2B (Figure 4A). We also over-expressed ΔNp63 in BEAS-2B (Figure 4A). 72 hours post transfection cells were exposed to increasing concentrations of CPPD for 48 hours (Fig 4B, 4C and S4A). While control cells were sensitive to CPPD and prone for DNA damage, indicated by ɣ-H2AX, overexpression of USP28 or ΔNp63 caused CPPD resistance, ɣ-H2AX clearance and increased cell survival (Figure 4B, 4C, 4D and S4A). Similar results were obtained in the USP28 mutant cell line Sk-Mes1. While the parental cell line expresses low levels of USP28 and is sensitive to CPPD, conditional overexpression of USP28 resulted in a higher tolerance of cisplatin (Figure S4B). It is noteworthy that exogenous USP28 led to an enhanced protein abundance of the USP28 target ΔNp63 in Sk-Mes1 (Figure S4B).

**Figure 4.**
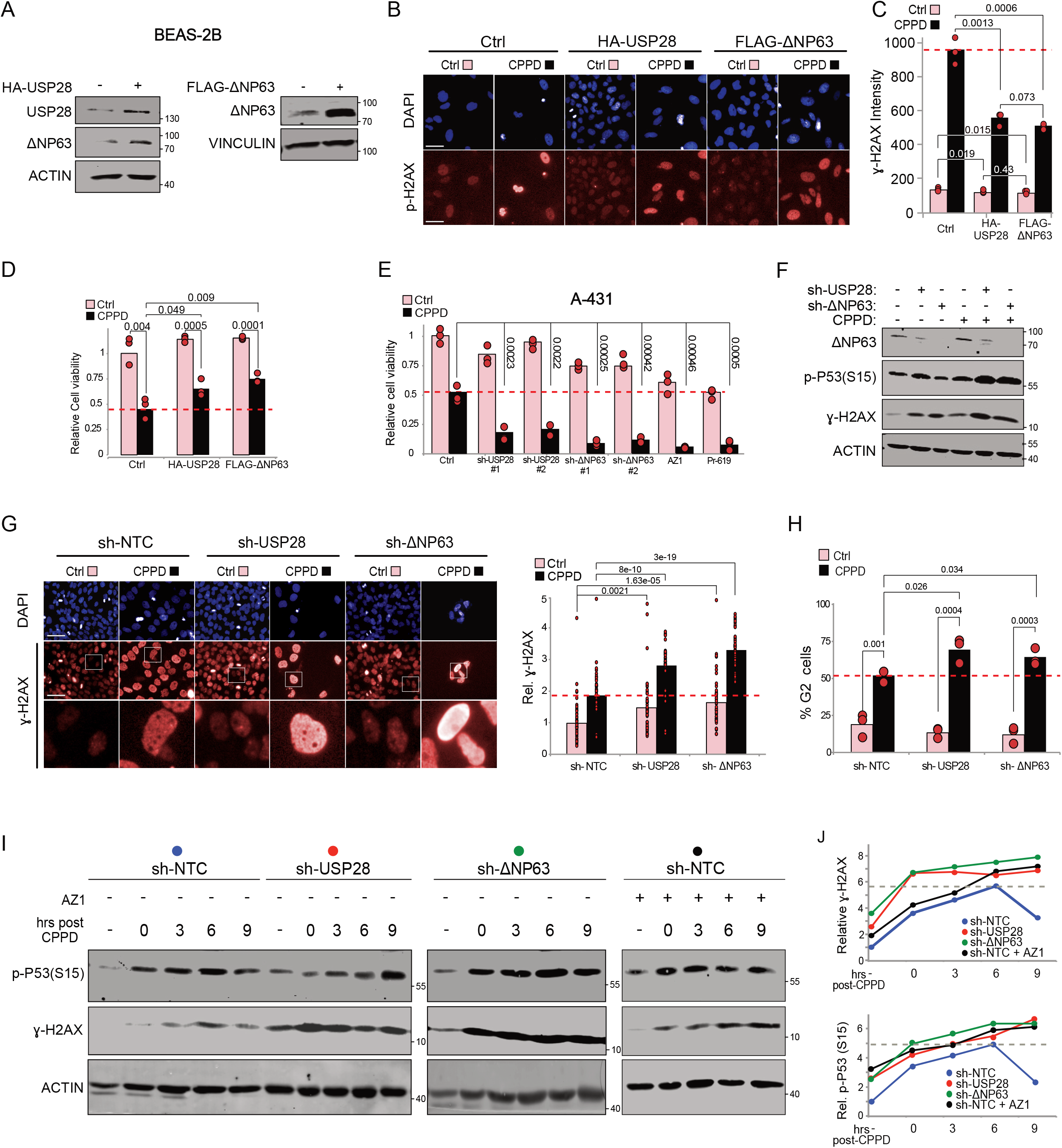
USP28-ΔNp63 axis is required for DDR upon cisplatin treatment and chemoresistance in SCC. A) Immunoblotting of BEAS-2B cells transiently transfected with either human USP28 or ΔNp63. Transfection of a GFP cDNA expressing plasmid served as control (-). ACTIN and VINCULIN served as loading control. n=3. B) Immunofluorescence staining against the DNA damage marker ɣ-H2AX in BEAS-2B cells transiently transfected with constructs from A) and exposed to 2.5μM CPPD for 48 hours. DAPI served as nuclear marker. n=3. Scale bar= 100μm C) Quantification of relative ɣ-H2AX fluorescence intensity in BEAS-2B from B). 15 fields per well (n=3) were quantified. P-values were calculated using two-tailed T-test statistical analysis. D) Quantification of relative cell survival in BEAS-2B from B). 15 fields per well (n=3) were quantified. P-values were calculated using two-tailed T-test statistical analysis. E) Quantification of relative cell survival of A431 control cells, either genetically depleted of USP28 or ΔNp63 by shRNA, treated with AZ-1 or the pan-DUB inhibitor PR-619, exposed to DMF or 5μM CPPD for 48 hours. n=3. F) Immunoblot of endogenous ΔNp63, phospho-serine 15 TP53, ɣ-H2AX in lentivirally transduced A431 cells (shRNA-control, shRNA USP28 or ΔNp63), exposed to DMF or 5μM CPPD for 24 hours. ACTIN served as loading control. n=3. G) Immunofluorescence staining against phospho-H2AX in lentivirally transduced A431 cells (shRNA-control, shRNA USP28#1 or ΔNp63#1) upon exposure to either DMF or 5μM CPPD for 48 hours. DAPI served as nuclear marker. n=3. Quantification of relative phospho-H2AX fluorescence intensity in A431 cells. n=50 cells. Scale bar= 200μm. P-values were calculated using two-tailed T-test statistical analysis. H) FACS-based cell cycle analysis and quantification of percentage of cells in G2 phase for lentivirally transduced A431 cells (shRNA-control, shRNA USP28#1 or ΔNp63#1) upon exposure to either DMF or 5μM CPPD for 48 hours. n=3. P-values were calculated using two-tailed T-test statistical analysis. I) Immunoblotting against endogenous phospho-P53 at serine 15 and phospho-H2AX in sh-control, sh-USP28#1, shΔNp63#1 or sh-control treated with AZ1. A431 cells treated with either DMF (-) or 5μM CPPD for 1 hour and collected at indicated time points after CPPD exposure. ACTIN served as loading control (n=3). See also Figure S4

Next, we depleted USP28 or its substrate ΔNp63 by shRNA knock down in A431 cells (Figure S4C) and treated these cells with either DMSO or CPPD for 72 hours and assessed cell viability (Figure 4E and S4C). While parental cells tolerated CPPD for 72 hours, cells depleted of USP28, ΔNp63 or treated with the inhibitors AZ1 or PR619 (pan-DUB inhibitor) showed reduced cell survival (Figure 4E). Analysing depleted cells by immunoblotting revealed that loss of USP28 or ΔNp63 already induced basal DNA damage, as seen by elevated levels of phosphorylated TP53 at serine 15 and ɣ-H2AX (Figure 4F). Upon exposure to CPPD, cells depleted for USP28 or ΔNp63 further increased the phosphorylation of TP53 and ɣ-H2AX, when compared to control cells (Figure 4F).

Since loss of USP28 and ΔNp63 resulted in an increased abundance of ɣ-H2AX, we wondered if knock down cells are already ‘primed’ for DNA damage. To address this question, we stained endogenous phosphor ɣ-H2AX by immunofluorescence in control and knock down cell lines, with and without treatment with CPPD (Figure 4G and S4D). While in non-stimulated cells ɣ-H2AX was hardly detectable – but increased upon treatment with CPPD – loss of USP28 or ΔNp63 resulted in increased ɣ-H2AX upon CPPD treatment (Figure S4D). As inhibition of USP28 resulted in an increased DNA damage response, we next considered if depletion of USP28 could affect cell cycle progression. Upon CPPD treatment, A431 cells accumulate during the G2 phase (Figure 4H and S4E). This was even more pronounced when USP28 or ΔNp63 was depleted (Figure 4H and S4E).

When cells were treated with 5μM AZ1 for 48 hours, a comparable accumulation of DNA damage to CPPD treatment was observed, as seen by the presence of TP53BP1 foci/nuclear bodies, demonstrating that inhibition of USP28 affects DNA integrity under basal conditions (Figure S4F). The extent of DNA damage was even further increased by combining AZ1 and CPPD for 48 hours, resulting in a significant overall increase in TP53BP1 foci/nuclear bodies positive cells (Figure S4F).

As genetic depletion of USP28 and ΔNp63 resulted in enhanced DNA damage upon CPPD treatment, we considered if USP28 and ΔNp63 are somehow involved in DNA damage recognition and/or clearance. We therefore performed a CPPD pulse-chase experiment in A431 control, knock down and AZ1 treated cells for 9 hours, and collected total protein at various time points to analyse the presence of phosphorylated TP53 and ɣ-H2AX by immunoblotting (Figure 4I). While in control cells TP53 and H2AX were phosphorylated rapidly upon CPPD treatment, the DNA damage response cleared within 9 hours post CPPD pulse. Cells under USP28 depletion or inhibition, however, showed elevated levels of phosphorylated TP53 and ɣ-H2AX, which further increased in the CPPD pulse, and these cells maintained elevated levels even after 9 hours post pulse (Figure 4I). Similar results were obtained by depleting ΔNp63; cells failed to clear the DNA damage 9 hours post CPPD pulse (Figure 4I).

These data demonstrate that USP28, potentially via ΔNp63, facilitates CPPD resistance and is involved in DNA damage clearance upon chemotherapy treatment.

### Disrupting the USP28-ΔNp63 Axis in SCC Affects Fanconi Anemia DDR Signature Genes

The Fanconi Anemia pathway is regulated by ΔNp63 and involved in mediating chemotherapy resistance(Bretz et al., 2016). Analyzing publicly available datasets revealed that in non-small cell lung cancer, SCC express elevated levels of USP28, ΔNp63 and FA pathway genes, compared to non-transformed or ADC samples (Figure 5A). Furthermore, SCC tumors frequently amplify ATR, while ATM is commonly downregulated or lost (Figure 5A). This is in stark contrast to adenocarcinomas, which upregulate ATM rather than ATR (Figure 5A). In contrast to SCC, ADC regulate the expression of FA pathway target via the E2F pathway(Hoskins et al., 2008). To assess if the USP28-ΔNp63 axis is a suitable target in re-establishing chemotherapy sensitivity, we analysed publicly available NSCLC datasets with regard to patient survival, chemotherapy and ΔNp63 expression (Figure 5B). Here, ΔNp63 is a strong indicator for chemotherapy failure in Non-Small Cell Lung Cancer (NSCLC), as the elevated expression of ΔNp63 correlates with poor survival under chemotherapy, while in patients receiving no chemotherapy, ΔNp63 has little prognostic value (Figure 5B and(Matin et al., 2013)). In lung SCC, ΔNp63 expression significantly correlates with FA, in particular in tumor tissue (Figure 5C). Similar correlations were identified for USP28 and FA, too (Figure 5D), indicating that USP28 contributes, via ΔNp63, to the regulation of FA in SCC. Cells depleted of ΔNp63 by two independent shRNA resulted in decreased FANCD2 protein abundance, even without CPPD or any other DNA damage agent (Figure S5A and S5B). CPPD pulse chase experiments in A431 cells demonstrated that FANCD2 is upregulated shortly after CPPD administration, reaching its peak expression around 6 hours post treatment (Figure S5C). Cells depleted of ΔNp63 fail to activate FANCD2 entirely during CPPD treatment (Figure S5C). The dependence of the FA pathway on ΔNp63 is hardwired into SCC. Loss of ΔNp63 resulted in a gross impairment of DDR and specially FA, as seen by RNA sequencing and mass spectrometry analysis of ΔNp63 knock down A431 cells (Figure S5D, S5E, S5F and S5G). Hence, ΔNp63 presents a vulnerability in SCC, which can be exploited by modulating its abundance.

**Figure 5.**
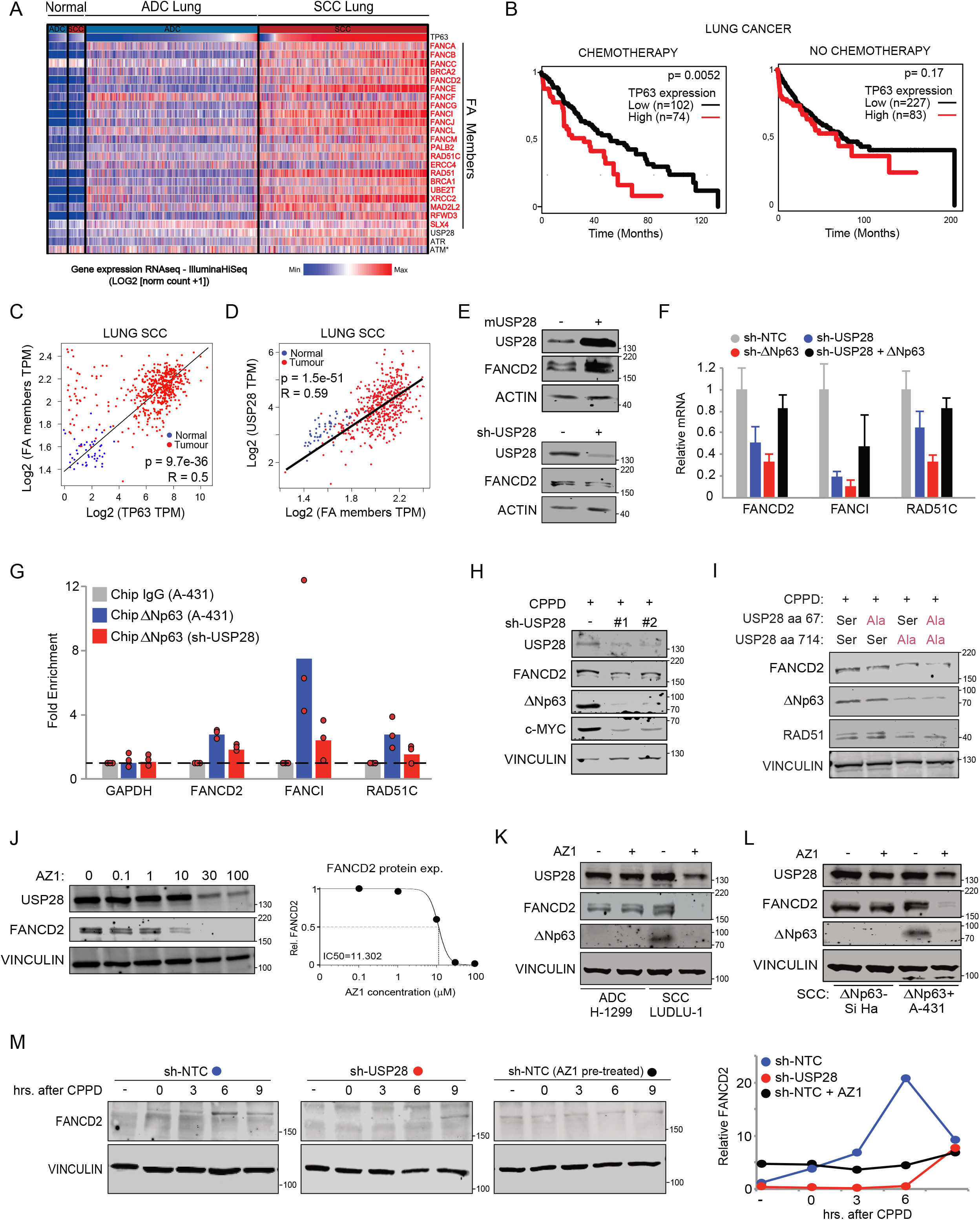
Deregulation of ΔNp63 impairs the Fanconi Anemia pathway in SCC. A) Publicly available gene expression analysis of TP63, Fanconi Anemia pathway genes, USP28, ATM and ATR in human non-transformed lung tissue, lung ADC and lung SCC. Generated with the online tool www.xena.ucsc.edu. Direct FA key members are highlighted in red. B) Publicly available patient survival data of NSCLC patients and stratified by relative expression of ΔNP63. Left panel= NSCLC patients treated with chemotherapy; Right panel= NSCLC patients non-treated with chemotherapy. Generated with the open source tool www.kmplot.com. C) Correlation between ΔNP63 and FA signature gene expression in human lung SCC and normal tissue. The diagonal line reflects a regression build on a linear model. R: Spearman’s correlation coefficient. Generated with the open source tool www.gepia2.cancer-pku.cn. D) Correlation between USP28 and FA signature gene expression in human lung SCC and normal tissue. The diagonal line reflects a regression build on a linear model. R: Spearman’s correlation coefficient. Generated with the open source tool www.gepia2.cancer-pku.cn. E) Immunoblot of USP28 and FANCD2 in A431 cells virally transduced with either doxycycline inducible overexpression of murine Usp28 or doxycycline inducible shRNA targeting USP28 for knock down. Cells were exposed to 1μ/ml for 96 hours prior to analysis. ACTIN serves as loading control. n=3. F) Quantitative RT-PCR of *FANCD2, FANCI* and *RAD51C* in A431 sh-NTC, sh-USP28#1, sh-ΔNp63#1 or sh-USP28#1 transfected with ΔNp63 cells normalised to ACTB. Quantitative graphic is represented as mean ± SD of three experiments (n= 3). G) RT-PCR of GAPDH, FANCD2, FANCI and RAD51C promotor regions upon Chromatin immuno-precipitation of either IgG or ΔNp63 in sh-NTC, sh-USP28#1 and sh-ΔNp63#1 A431 cell lines. Normalised to IgG. Quantitative graphic is represented as mean ± SD of three experiments (n= 3). H) Immunoblot of endogenous USP28, FANCD2, ΔNP63 and c-MYC in A431 cells harbouring two independent inducible shRNA targeting USP28. Cells were exposed to 1μg/ml for 72 hours prior to analysis, followed by 24 hours of 1μg/ml doxycycline and 5μM CPPD co-treatment. VINCULIN serves as loading control. n=3. I) Immunoblot of endogenous FANCD2, ΔNP63 and RAD51 in control, S67A, S714A and S67A+S714A mutant A431 cells treated with 5μM CPPD for 6 hours. VINCULIN serves as loading control. n=3 J) Immunoblot of endogenous USP28 and FANCD2 in A431 cells treated for 24 h with either DMSO or indicated concentrations of AZ1. VINCULIN served as loading control. FANCD2 half-maximal inhibitory protein abundance (IC_50_) was calculated. K) Immunoblot of USP28, FANCD2 and ΔNp63 in control or AZ1 (15μM) treated lung cells H1299 (ADC) and LUDLU-1 (SCC). VINCULIN served as loading control. n=3. L) Immunoblot of USP28, FANCD2 and ΔNp63 in cervix SiHa (ΔNp63-) and vulva A431 (ΔNp63+) cells treated with DMSO or AZ1 (15μM). VINCULIN served as loading control. n=3. M) Immunoblot of FANCD2 in CPPD pulse chase experiment (5μM, 1 hour exposure, followed by washout) of control (sh-NTC), sh-USP28#1 or sh-NTC+AZ1 in A431 cells. Numbers indicate hours post CPPD treatment. VINCULIN served as loading control. n=3. Graph represents band intensity of FANCD2 in cells assessed by immunoblotting. See also Figure S6 and S7

Next, we conditionally over-expressed murine USP28 in A431 cells (Figure 5E). Upon doxycycline treatment for 96hours, cells showed enhanced protein abundance for USP28 and FANCD2 (Figure 5E). Conversely, conditional depletion of USP28 by an shRNA resulted in decreased FANCD2 protein abundance and mRNA expression (Figure 5E). Notably, mono-ubiquitylation of FANCD2 was not affected. This observation indicates that USP28 contributes to FA pathway regulation already under basal conditions by affecting ΔNp63 protein stability. Overexpression of ΔNp63 partially restored the expression of FANCD2, FANCI and RAD51C in USP28 depleted cells (Figure S5B and 5F). This was further supported by chromatin immunoprecipitation of ΔNp63 in control and USP28 depleted A431 cells (Figure 5G). While ΔNp63 was bound to the promoters of FANCD2, FANCI and RAD51C, loss of USP28 resulted in a significant reduction in ΔNp63 binding to its cognate recognition sites in these promoters (Figure 5G). SCC cells require USP28 to respond to DNA damage stress, as USP28 knock down A431 cells fail to stabilise and upregulate FANCD2, ΔNp63 or c-MYC upon CPPD treatment (Figure 5H). Similar effects were observed in the ATR SQ/TQ motive mutant A431 cells. These cells showed diminished protein levels of ΔNp63, FANCD2 and RAD51, in particular in the S714A and double mutant (Figure 5I).

As USP28 can be targeted by the pharmacologic inhibitor AZ1, we exposed A431 cells to increasing concentrations of the small molecule inhibitor (Figure 5J). Immunoblotting against FANCD2 revealed that inactivation of USP28 resulted in depletion of FANCD2 (FIG 5J). To investigate if this effect is SCC specific, we compared the expression of FANCD2 upon treatment with AZ1 in a lung adenocarcinoma cell line, NCI-H1299, versus an SCC cell line, LUDLU-1 (Figure 5K). Furthermore, to identify if the effect of USP28 inhibition is via ΔNp63, we compared the two SCC lines Si Ha (ΔNp63^negative^) and A431 (ΔNp63^positive^) (Figure 5L). While FANCD2 was detectable in all tested cell lines, only ΔNp63 expressing cells lost FANCD2 expression upon exposure to AZ1, along with ΔNp63 itself (Figure 5K and 5L). This effect was also observed in various SCC cell lines; exposure to AZ1 reduced FANCD2 protein abundance in Detroit 562 (HNSC), Ludlu-1 (LUSC) and Ca Ski (CESC) cells as well (Figure S5H).

The impairment of the FA pathway became obvious by performing CPPD pulse chase experiments in either USP28 knock down or AZ1-treated A431 cells (Figure 5M). In control cells, upon exposure to CPPD, FANCD2 was rapidly upregulated and mono-ubiquitylated. By immunoblotting the maximum activity of FANCD2, it was detectable up to 6 hours after CPPD treatment. USP28 depleted cells, or cells exposed to AZ1, failed to upregulate and activate FANCD2 at all (Figure 5M).

### Pharmacologic Inhibition of USP28 Re-sensitizes SCC Cells to Chemotherapy

By treating several human cancer cell lines with 2μM CPPD for 96 hours, we could observe that SCC, and in particular ΔNp63 expressing cells, tolerated CPPD better then adenocarcinoma cell lines (Figure S6A and B).

If ΔNp63 mediates CPPD resistance, and USP28 regulates ΔNp63 protein abundance, treatment with AZ1 could synergize with CPPD. To test this hypothesis, we exposed human SCC and, where applicable, same tissue ADC cells to various amounts of AZ1 in the presence of CPPD, thereby aiming to identify potential additive or synergistic effects (Figure 6A, 6B, S6C, S6D, S6E and S6F). Cells were seeded in 384 well plates and treated for 48 hours, followed by nuclear staining with DAPI and immunofluorescence staining against the DNA damage marker ɣ-H2AX (Figure 6A and S6C). While the ΔNp63 negative cell lines NCI-H1299 and HeLa showed no additive nor synergistic effect in cell viability upon co-treatment with CPPD and AZ-1, the ΔNp63 expressing SCC cell lines A431, Detroit 562, LUDLU-1 and Ca Ski showed synergistic effects when combining both compounds, as measured by LOEWE-synergism (Figure 6B and S6D). As already observed in A431 cells, SCC cells impaired for USP28 extended DNA damage signalling and failed to repair DNA damage over time, which was seen by ɣ-H2AX expression in cells exposed to CPPD, AZ1 or the combination of both (Figure 6A and S6C). In ADC cells, however, AZ-1 either impaired or accelerated ɣ-H2AX foci clearance, and the combination of CPPD and AZ-1 revealed a reduction in ɣ-H2AX staining intensity (Figure 6A and S6C).

**Figure 6.**
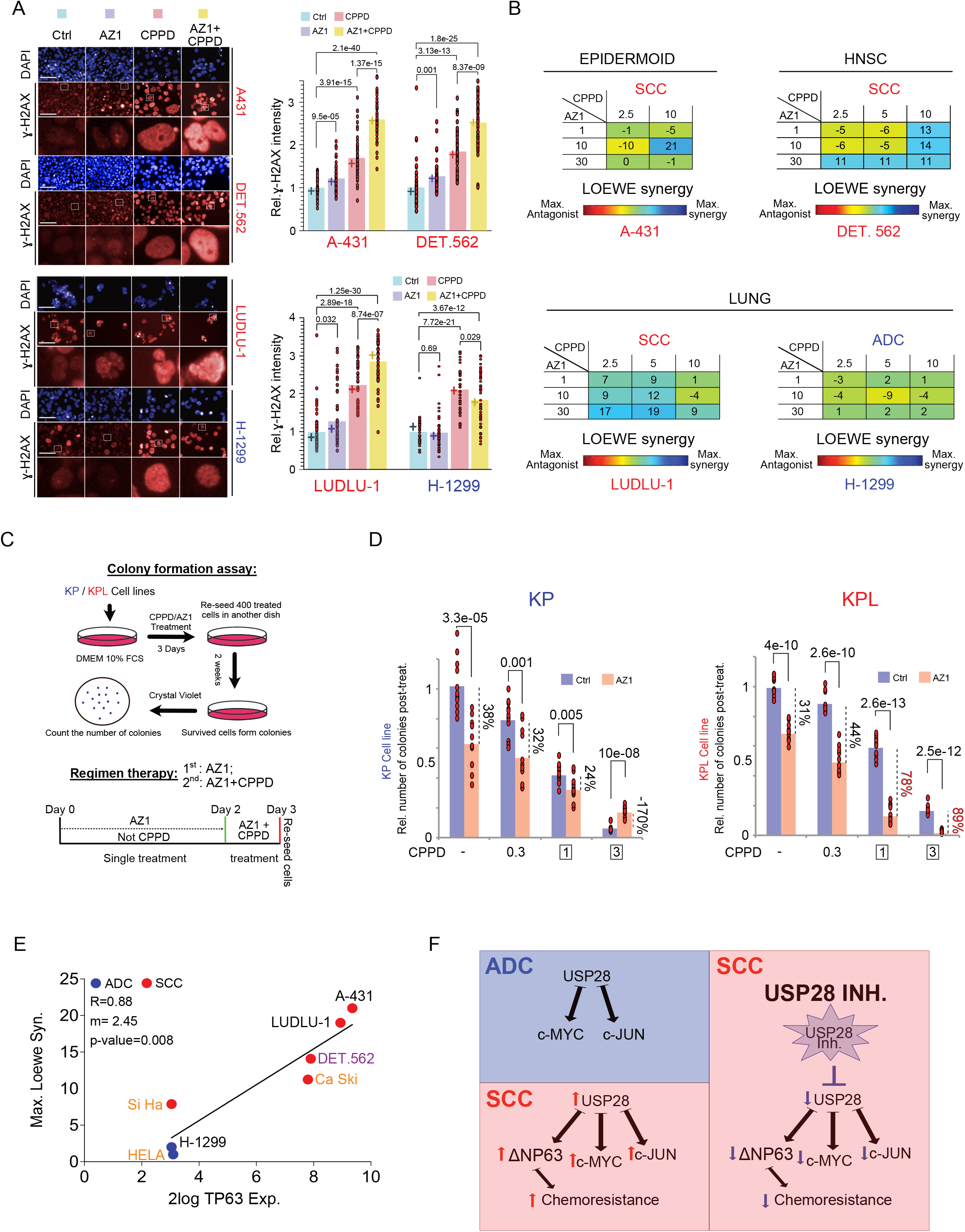
Pharmacologic inhibition of USP28 re-sensitizes SCC cells to chemotherapy. A) Immunofluorescence staining of phospho-H2AX in A431, Detroit 562, H-1299 and LUDLU-1 cells treated with DMSO+DMF (Ctrl), 15μM AZ1, 5μM CPPD or 15μM AZ1+5μM CPPD for 48 hours. DAPI served as nuclear marker. Relative quantification of the phospho-H2AX staining intensity was measured for the different treatment exposures. n= 50 cells. Scale bar= 200μm. P-values were calculated using two-tailed T-test statistical analysis. Red= SCC cell line; Blue= ADC cell line. B) LOEWE synergism score of CPPD and AZ1 in A431, Detroit 562, H-1299 and LUDLU-1 cell lines. Cells were exposed to the Indicated concentrations (μM) for 48 hours. DAPI was used to assess total cell numbers. Red= SCC cell line; Blue= ADC cell line. C) Schematic representation of the colony formation assay in KP / KPL murine cell lines obtained from in lung vivo tumors. Red= SCC cell line; Blue= ADC cell line. D) Relative number of colonies after treatment with either DMSO or 15μM AZ1 and exposure to CPPD at indicated concentrations in KP and KPL cell lines. Experiments was performed as described in Figure 5C. n=11. P-values were calculated using two-tailed T-test statistical analysis. E) Spearman correlation of TP63 mRNA expression and maximum LOEWE synergism observed. The diagonal line reflects a regression build on a linear model. R: Spearman’s correlation coefficient, m: slope of the linear regression mode. F) Model of USP28 action in chemotherapy resistance in SCC versus ADC. See also Figure S5

To identify if ΔNp63, and not the tumor type, is mediating chemoresistance, next treated an ADC cell line that expresses low levels of ΔNp63, A549, as well an SCC cell line that lost expression of ΔNp63, Si-Ha with CPPD, AZ1 or the combination of both compounds (Figure S6C, S6E and S6F). After treatment with both compounds, A549 showed an increased ɣ-H2AX expression and less cell survival, when compared to CPPD or AZ1 alone, thereby behaving similar to SCC cell lines (Figure S6C and S6E). Si-Ha, despite being of SCC origin, showed no additive nor synergistic effect with combinatorial treatment, and the exposure to AZ1 resulted in reduced levels of γ-H2AX, compared to CPPD alone (Figure S6C and S6F).

As ADC and SCC cells showed differential responses to AZ1/CPPD combinatorial treatment, we wondered if ADC cells could become resistant to CPPD. To address this question we used our previously established murine lung cancer cell lines, KP (ADC, *Kras^G12D^, Trp53^Δ^*) and KPL (ΔNp63p^ositive^ SCC, *Kras^G12D^, Trp53^Δ^, Lkb1^Δ^*) (Figure S6G)(Prieto-Garcia et al., 2020b). Cells were seeded and cultured in the presence of AZ1 for 2 days, followed by 1 day with AZ1/CPPD combinatorial treatment. Surviving cells were re-seeded and cultured for 2 weeks in either AZ1, CPPD or both. Finally, the remaining colonies were stained by Crystal violet and counted (Figure 6C and 6D). When cells were exposed to CPPD alone, KPL cells tolerated CPPD better then KP cells (Figure S6G), in line with human cell line responses to CPPD. Co-exposure with AZ1, however, sensitized KPL cells to CPPD and led to a significant increase in cell death at 1μM CPPD, when compared to KP (Figure 6D). It is noteworthy that in high concentrations of CPPD (3μM), the addition of AZ1 to the ADC cell line KP resulted in an increased appearance of resistant clones, while the SCC line KPL succumbed to the treatment (Figure 6D).

Not only does treatment with AZ1 sensitize SCC cells to CPPD, but it synergises with additional DNA damaging agents, such as Oxaliplatin or 5-FU, as seen by viability assays in A431 cells (Figure S6I).

Hence, inhibition of USP28 synergizes with CPPD predominantly in cells expressing ΔNp63, while in ADC or SCC cells lacking ΔNp63 expression, no cooperation between CPPD and USP28 inhibition could be observed (Loewe synergy, Spearman R= 0.88, Figure 6E and 6F). Furthermore, our data suggest that in ADC cells inhibition of USP28 by AZ1 could result in cell cycle stop and thereby support a more efficient DNA damage clearance and hence foster the rise of chemotherapy resistant cancer cells.

### Inhibition of USP28 activity deregulates FA-DDR signalling in vivo and sensitizes tumors to CPPD treatment in ex vivo organotypic lung SCC tumor slice cultures

To assess if the effects of USP28 on FA are preserved *in vivo*, we analyzed the expression of USP28, ΔNp63 and DDR markers in a murine model of lung SCC. We utilized a previously characterized constitutive CRISPR/Cas9 expressing mouse strain, in combination with AAV virions, for tumor induction and depletion of USP28 in the lung (KPL (<u>K</u>RasG12D:T<u>p</u>53^A^:<u>L</u>kb1^A^) versus KPLU (<u>K</u>RasG12D:T<u>p</u>53^A^:<u>L</u>kb1^A^:<u>U</u>sp28^A^) (Figure S7A and S7B)(Prieto-Garcia et al., 2020a). Immuno-histochemical analysis of lung sections of KPL mice showed that USP28 was readily detectable, and the DDR markers TP53BP1, ɣ-H2AX and the FA effector FANCD2 were expressed (Figure S7C). In KPLU tumors the DDR markers were strongly upregulated, while FANCD2 was not detectable (Figure S7C). Western blot analysis of primary tumor material comparing KPL and KPLU showed the depletion of USP28 and loss of ΔNp63, as previously described (Figure S7D)(Prieto-Garcia et al., 2020a). In KPLU tumors, the overall protein abundance of the FA pathway proteins FANCD2 and FANCI were significantly reduced, when compared to USP28-proficient tumor samples (Figure S7D), demonstrating that the USP28-ΔNp63 axis is required for FA expression *in vivo*.

As systemic inhibition of USP28 by the small molecule inhibitor AZ1 is well tolerated in mice(Prieto-Garcia et al., 2020a), we wondered if deregulated DDR via inhibition of the FA pathway could be observed in AZ1-treated animals, thereby contributing to the observed anti tumor effect (Figure 7A and B). Immuno-histochemical analysis of tumor-bearing lungs from murine SCC transplant animals (KPL and Figure S6G) revealed that in control treated animals USP28 and its substrate ΔNp63 were detectable, along with the ΔNp63 target FANCD2 (Figure 7C and S7E). DNA damage markers, such as TP53BP1 and ɣ-H2AX, were only weakly expressed (Figure 7C). The amount of detectable TP53BP1 and ɣ-H2AX could reflect basal DDR activity due to ongoing transcription/replication in tumor cells(Fernandez-Vidal et al., 2017; Kotsantis et al., 2016).Tumor-bearing animals exposed to AZ1, however, showed reduced detectability of USP28 and ΔNp63, as previously described(Prieto-Garcia et al., 2020a). Compared to control animals, FANCD2 was only weakly expressed and the DDR markers TP53BP1 and ɣ-H2AX upregulated (Figure 7C and 7D). This was further confirmed by tumor tissue explants, following immunoblotting (Figure 7E), revealing a significant reduction in USP28, ΔNp63 and FANCD2 upon treatment with AZ1 (Figure 7E).

**Figure 7.**
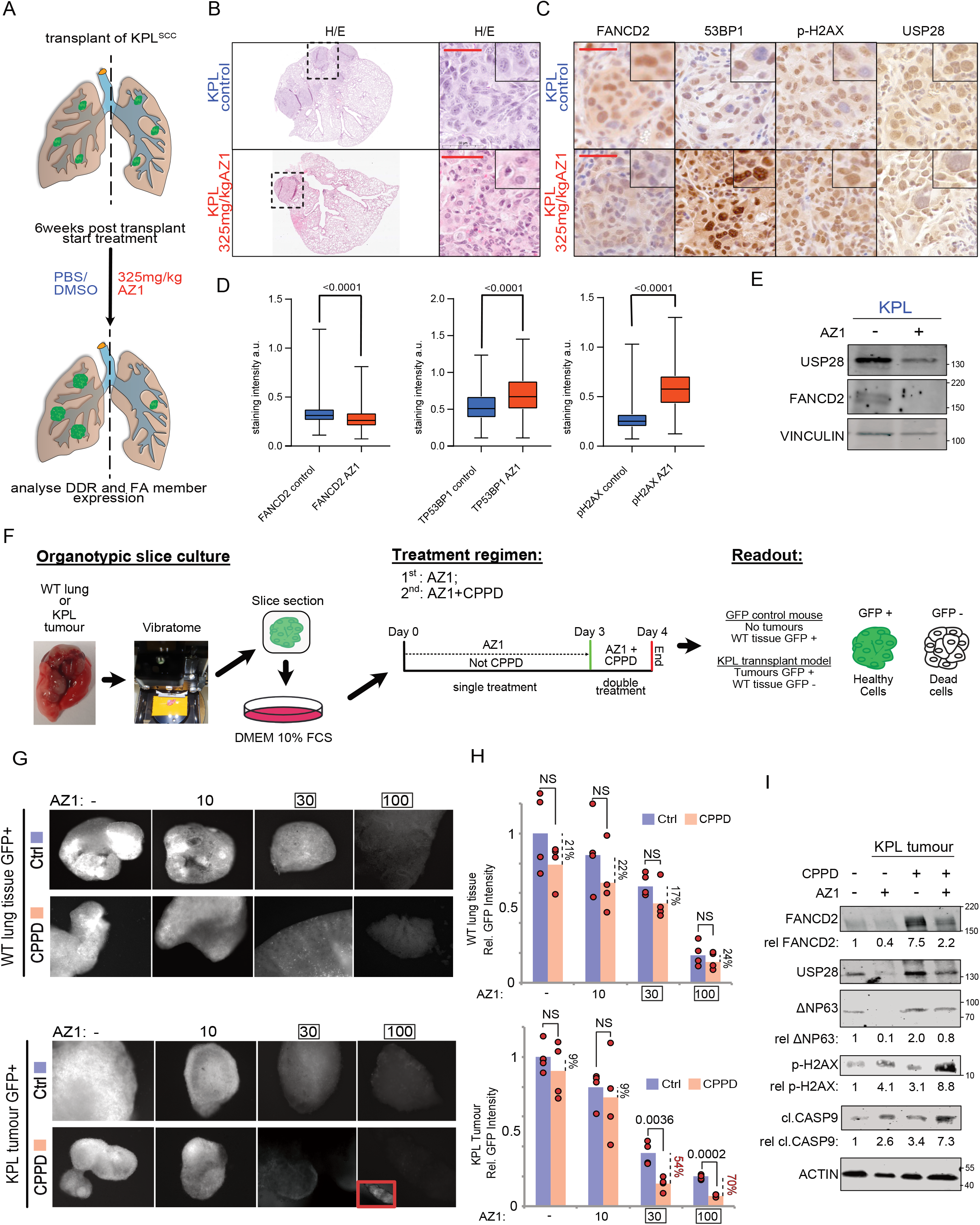
Inhibition of USP28 activity deregulates FA-DDR signalling in vivo and sensitizes tumors to CPPD treatment in ex vivo organotypic lung SCC tumor slice cultures by de-activating FA. A) Schematic diagram of isogenic transplant experiment of of *p53^Δ^; Lkb1^Δ^; KRas^G12D^(KPL*) cells, followed by treatment with either vehicle (PBS/DMSO/Tween) or AZ1 (375mg/kg), for a total of 18 days. B) Representative images of sections from KPL and KPLU mice stained with haematoxiline and eosine. Inlay shows higher magnification. Scale bar = 50μm C) Immunohistochemistry of FANCD2, TP53BP1, phospho-H2AX and USP28 in KPL transplant tumors post treatment with either vehicle (PBS/DMSO/Tween) or AZ1 (375mg/kg), for a total of 18 days. Scale bar = 50μm D) Quantification of relative immunohistochemical staining intensity of FANCD2, γ-H2AX and TP53BP1 in vehicle or AZ1 treated mice. Statistical analysis was performed using unpaired t test. P<0.0001. Images were quantified using QuPath (version0.2.8). Boxplots were generated using Graphpad Prism8. a.u.= Arbitrary units. n^FABCD2^=6200 control/3616 AZ1; n^ɣ-H2AX^ =17075control/9140 AZ1; n^TP53BP1^=18831 control/8586 AZ1. P-values were calculated using two-tailed T-test statistical analysis. E) Immunoblot of endogenous USP28 and FANCD2 from KPL tumors post treatment with either vehicle (PBS/DMSO/Tween) or AZ1 (375mg/kg), for a total of 18 days. VINCULIN served as loading control. n = 3. F) Schematic model of ex-vivo organotypic slice culture model of lung cancer (see Material and Methods section for details) and experimental design. G) Immunofluorescence of WT GFP^+^ lung cells within the organotypic slice culture after 72 hours of indicated AZ1 concentrations and 24 hours of co-treatment with 5μM CPPD. Quantification of relative GFP^+^ signal intensity of organotypic slice cultures. n=4. P-values were calculated using two-tailed T-test statistical analysis. H) Immunofluorescence of KPL (*p53*^Δ^; *Lkb1*^Δ^; *KRas^G12D^*) GFP^+^ lung tumor cells within the organotypic slice culture after 72 hours of indicated AZ1 concentrations and 24 hours of co-treatment with 5μM CPPD. Quantification of relative GFP^+^ signal intensity of organotypic slice cultures. n=4. P-values were calculated using twotailed T-test statistical analysis. I) Immunoblot of KPL (*p53*^Δ^; *Lkb1*^Δ^; *KRas^G12D^*) GFP^+^ lung tumor cells within the organotypic slice culture after 72 hours of indicated AZ1 concentrations and 24 hours of co-treatment with 5μM CPPD. Quantification of relative protein signal using Actin for normalisation. n=4.

We next tested different treatment regimes in A431 cells for using AZ1 and CPPD (Figure S7F, S7G and S7H), comprising single compound treatment, AZ-1/CPPD co-treatment, AZ-1 pre-treatment followed by CPPD and AZ-1 pre-treatment, followed by AZ-1/CPPD co-treatment (Figure S7F). Upon exposure to the various treatment regimes, cells were fixed, followed by immunofluorescence staining against TP53BP1 and quantification of the total number of cells and cells with >25 TP53BP1 foci/field of view (Figure S7F and G). AZ-1 showed the weakest DDR staining intensity, followed by cells exposed to CPPD alone (Figure S7F and G). Either simultaneous or consecutive treatment of cells with AZ-1 and CPPD resulted in similar cell survival and TP53BP1 foci formation (Figure S7F and G). Pre-treatment with AZ-1 followed by co-exposure to the small molecule inhibitor and the chemotherapeutic agents resulted in a strong upregulation of TP53BP1, including an even distribution within the nucleus, and led to an enhanced cell death in A431 cells (Figure S7F and G).

To investigate a potential therapeutic synergism between AZ-1 and CPPD in a multicellular system, we decided to employ the *ex vivo* organotypic lung slice culture (Figure 7F). Here, isogenic murine SCC cells (KPL) were orthotopically re-transplanted in immune-competent C57BL6/J mice where they engrafted and formed a tumor (Figure 7F). 8 weeks’ post-transplant, mice were sacrificed and the tumor-bearing lungs explanted, following live tissue sectioning with a vibratome (Figure 7F). As a control, we used a lung slice culture from a wild type *C57BL6/J-Rosa26^Sor-CAGG-Cas9-IRES-eGFP^* mouse. Slices containing tumor (GFP^+^) or wild type tissue (GFP^+^) were cultured in standard medium (DMEM, 10%FCS) and exposed to small molecule inhibitor AZ1 (0-100 μM AZ-1) and CPPD (5μM) according to treatment regime (Figure S7F). While AZ-1 treatment alone had an anti-proliferative, pro apoptotic effect on GFP+ KPL cells, co-treatment of the organotypic slice culture with AZ-1 and CPPD significantly reduced the amount of detectable, viable tumor cells, at around ~30μM AZ1/ 5μM CPPD (Figure 7G and H). In contrast, wild type lung slice cultures exposed to the same treatment regime tolerated the dosages significantly better and showed weaker responses, as measured by GFP intensity (Figure 7G and H). Immunoblotting of tissue samples from tumor-bearing organotypic slice cultures post treatment revealed that AZ-1 single treatment efficiently depleted FANCD2 protein abundance, but upon co-treatment with CPPD, lung slices lost FANCD2 and significantly upregulated ɣ-H2AX and pro-apototic signaling, as seen by cleaved Caspase 9 (Figure 7I).

These data show that inhibition of USP28 specifically affect tumor cell growth by interfering with the DDR pathway and that priming of SCC cells via AZ-1 potentiates the therapeutic efficacy of AZ-1/CPPD co-treatment *in vitro* and *ex vivo*.

## DISCUSSION

### USP28 in DNA damage signalling

Maintaining chromosomal stability by minimizing the accumulation of DNA damage is a prerequisite for cells to survive(Hanahan and Weinberg, 2011; Jackson and Bartek, 2009). Fast proliferative cells in particular require efficient mechanisms to cope with consistent single/double strand breaks and DNA damage, and therefore depend on the ability to resolve transcription-replication conflicts(Hamperl and Cimprich, 2016). As a consequence, several therapeutic strategies aim at inflicting DNA damage to overwhelm the DNA damage repair machinery in tumor cells, as these cells, in contrast to non-transformed cells, frequently harbor mutations in check point genes and fail to halt the cell cycle to initiate the repair of damaged DNA(Bhattacharya and Asaithamby, 2017; Khanna, 2015; Medema and Macurek, 2012; Nikolaev and Yang, 2017). Previous reports demonstrated that deubiquitylating enzymes are involved in the DNA damage response pathway, including USP28, which is recruited to sites of ionizing radiation induced DNA damage, where it interacts with TP53BP1 and ATM(Kee and Huang, 2016; Pinto-Fernandez and Kessler, 2016; Zhang et al., 2006). This interaction results in the phosphorylation of the DUB and leads to its dissociation from its interacting E3 ligases(Popov et al., 2007a; Zhang et al., 2006; Zhang et al., 2020). The overall role of USP28 in DNA damage signalling, however, was unclear(Knobel et al., 2014).

In this study we identified that USP28 is recruited to sites of cisplatin-induced DNA damage in SCC cells, along with other DNA damage readers. Upon exposure to cisplatin, USP28 interacts with ATR and ATM, while ATR-dependent phosphorylation of USP28 represents the predominant DNA damage induced modification of USP28. This is important, as the cisplatin-induced interaction with ATR, and the subsequent phosphorylation of USP28, results in an increase in USP28 enzymatic activity. As a consequence, the ability of USP28 to stabilize substrates, such as MYC and ΔNp63, is enhanced during cisplatin treatment. Inhibition of the DNA damage kinase ATR by VE-821 blocked the phosphorylation of USP28, which resulted in its inactivation, as seen by ubiquitin suicide probe assays. Furthermore, indirect inhibition of USP28 activity by blocking ATR function resulted in the degradation of USP28, in line with previous reports which identified that DUB protein stability is linked to their activity(de Bie and Ciechanover, 2011; Wang et al., 2018). With the advent of small molecule inhibitors, targeting deregulated protein stability by inhibiting deubiquitylases, or the proteasome, became a feasible strategy(An et al., 2017; Colland et al., 2009; Fan et al., 2013; Lamberto et al., 2017; Wrigley et al., 2017). Loss of USP28, upon genetic or pharmacologic inhibition with the small molecule inhibitor AZ1, reduced the abundance of DNA damage signature proteins in SCC. Intriguingly, this preferentially affected specific DNA damage repair mechanisms.

In SCC, ΔNp63 is an essential factor regulating chemoresistance as it drives the expression of DNA damage genes(Ratovitski, 2014; Sen et al., 2011). Controlling the protein abundance of ΔNp63 by USP28, therefore, directly affects chemosensitivity. While SCC cells rapidly cleared cisplatin-induced DNA damage, loss/inhibition of USP28 or ΔNp63 resulted in persistent activated DNA damage signalling and the inability of cells to clear the damage markers ɣ-H2AX, along with prolonged serine 15 phosphorylation of TP53. The inability of SCC to clear the damage signature indicates that the repair machinery is impaired, which in turn can be leveraged to enhance and re-establish a cisplatin response in otherwise chemoresistant human SCC cell lines. ΔNp63 expressing SCC therefore showed a significant synergism between AZ1 and cisplatin. AZ1 alone led to ɣ-H2AX foci formation, which was further enhanced upon co-treatment with cisplatin, resulting in reduced overall survival in a dose-dependent fashion.

It is worth noting that human adenocarcinoma cell lines, and the SCC line SiHa, which does not express ΔNp63, behaved differently. The inhibition of USP28 here led to a reduction of ɣ-H2AX, and co-treatment with cisplatin had no additive nor synergistic effect. This is in line with previous reports, where loss of USP28 in a NSCLC adenocarcinoma cell line H460 mediated resistance to ionising radiation(Zhang et al., 2006). Similar observations were made in other tumor entities, where loss of USP28 induced treatment resistance(Saei et al., 2018).

Modulation of USP28 protein abundance or enzymatic activity alone already affected tumor burden and tumor maintenance *in vivo(Prieto-Garcia et al., 2020b).* Here, loss or impairment of USP28 resulted in a significant increase in DNA damage marker abundance, namely ɣ-H2AX and TP53BP1, while the Fanconi Anemia pathway was reduced. By employing an *ex vivo* organotypic slice culture system, we could further demonstrate that the combination of USP28 inhibition and DNA damage, induced by cisplatin, resulted in tumor shrinkage, while wild type tissue tolerated the treatment.

USP28 behaves like an NOA (non-oncogene addiction) gene, such as wild type cells tolerate its inactivation, while tumor cells, and in particular SCC, depend on its expression(Diefenbacher and Orian, 2017; Diefenbacher et al., 2014). Animals, and in particular tumor cells, undergo DDR stress upon inhibition of USP28 via the small molecule AZ1. Hence, tumor cells are primed for DNA damage by USP28 inhibition and are therefore more susceptible to DNA damaging agents, such as CPPD.

### USP28, the double edged sword

Targeting USP28, either genetically or pharmacologically, resulted in reduced proliferation in *in cellulo* assays and reduction in tumor burden *in vivo* (Prieto-Garcia et al., 2020a). In lungs, both major NSCLC sub-entities, ADC and SCC, required USP28 for proliferation, while SCC were dependent on USP28 for tumor induction(Prieto-Garcia et al., 2020a).

We therefore decided to combine USP28 inhibition with CPPD to target tumor cells. While using AZ-1 to inhibit USP28 in ADC and SCC cells, we identified several striking differences among the two different NSCLC entities. In ADC, USP28 is required to stabilize the onco-proteins JUN, MYC and NOTCH, and cells depend on these factors to facilitate proliferation *in vitro.* SCC, however, require ΔNp63 and NOTCH as major drivers of proliferation. Upon exposure to the USP25/28 inhibitor AZ1, in contrast to SCC, ADC in general tolerated higher doses of AZ1 and showed a strong reduction in ɣ-H2AX-staining intensity. Furthermore, ADC exposed to the combinatorial treatment with AZ1 and CPPD decreased the DNA damage marker ɣ-H2AX when compared to CPPD treatment alone. In contrast to SCC, ADC did not alter the expression of the FA pathway upon inhibition of USP28. The pathway mediating chemotherapy resistance is therefore still active in ADC, while SCC potentially fail to form the FA complex and so rely on alternative and error-prone DDR pathways(Bhattacharjee and Nandi, 2017; Sumpter and Levine, 2017). This results in the accumulation of DNA damage and subsequent cellular death. Loss or inhibition of USP28 in SCC results in the loss of ΔNp63, and consequently in the loss of the expression of its target genes, including FA pathway members such as FANCD2(Bretz et al., 2016; Hoskins et al., 2008).

Based on our findings, USP28 presents a promising therapeutic target, in combination with DNA-damaging agents such as CPPD, in SCC; while in ADC, due to the ΔNp63-independent expression of FA proteins by the E2F protein family(Hoskins et al., 2008), targeting of USP28 could potentially have adverse effects and even support the establishment of therapeutic resistance.

Overall, our results show that targeting the USP28-ΔNp63 axis in SCC tones down the Fanconi Anemia-DNA damage response pathway, thereby sensitizing SCC cells to cisplatin treatment.

## MATERIAL AND METHODS

### Tissue culture and reagents

A-431, Beas-2B, SiHa, Ca SKI, DETROIT 562, HEK-293T, NCI-H1299. cell lines were obtained from ATCC or ECACC. The human lung cancer cell line LUDLU-1^adh^ was described previously (Prieto-Garcia et al., 2020a). A-431, Beas-2B, SiHa, Ca SKI, DETROIT 562 and HEK-293T cells were cultured in DMEM (Gibco) supplemented with 10% fetal bovine serum (FCS)/ 1% Pen-Strep. LUDLU-1^adh^, NCI-H1299, CALU 1, SK-MES1, and H23 cells were cultured in RPMI 1640 (Gibco) supplemented with 10% FCS/ 1% GlutaMAX/ 1% Pen Strep. Cell lines were authenticated by STR profiling. Cells were routinely tested for mycoplasma via PCR.

Except when a different concentration was expressly indicated, the reagents were dissolved in Dimethyl sulfoxide (DMSO) or Dimethylformamide (DMF) and added to the cells at the following concentrations: Cisplatin (CPPD; 5μM; dissolved in DMF), doxycycline (DOX; 1μg/ml), Tandem ubiquitin binding entity (TUBE; 100 μg/ml), KU55933 (15 μM; dissolved in DMSO) and VE 821 (2.5 μM; dissolved in DMSO).

### DNA transfection and infection

DNA transfection was performed adding a mix of 2.5μg plasmid DNA, 200μl serum free medium and 5μl PEI to the cells seeded in a 6-well plate (60% confluence), after 6h incubation at 37°C the medium was changed to full supplemented medium and finally, cells were collected after 48 hours for experimental purposes. For viral infection, AAVs or Lentiviruses (MOI=10) were added to the medium in the presence of polybrene (5μg/ml) and incubating at 37°C for 4 days. The selection of infected cells was performed with 2,5 μg/ml Puromycin for 72h, 250μg/ml Neomycin for 2 weeks or FACS-sorting RFP/GFP positive cells (FACS Canto II BD).

### Primary murine lung cancer cell lines and colony formation assay

Primary lung cancer cell lines were obtained from 12 weeks old mice as previously described (Prieto-Garcia et al., 2020a). At endpoint of experiment, mice were sacrificed and lung tumors isolated. Tissue was digested in Collagenase I (100U/ml in PBS for 30 minutes at 37C and after stopping the reaction with FCS, the mixture was centrifuged and re-suspended in DMEM (Gibco) supplemented with 10% fetal bovine serum (FCS) and 1% Pen-Strep. Fibroblasts were counter-selected by selective trypsinisation and homogeneous cell clusters were clonally expanded. All clones have been characterized and classified according to markers as adenocarcinoma (KP cell lines) or squamous cell carcinoma (KPL cell line).

For colony formation assay, murine cells were treated at indicated concentrations of CPPD/AZ1 (Fig 5G and S5H) for 3 days. After exposure, 400 cells were re-seeded in a new 10cm plate and maintained in DMEM supplemented with 10% fetal bovine serum (FCS) and 1% Pen-Strep for 14 days. Number of healthy KP/KPL colonies was quantified manually upon staining with 0.5% Crystal violet.

### RT-PCR and CHIP-QPCR

RNA was isolated with Peq GOLD Trifast (Peqlab), as indicated in the manufacturer’s instructions. RNA was reverse transcribed into cDNA using random hexanucleotide primers and M-MLV enzyme (Promega). ChIP experiments were performed using 20μg anti-ΔNp63 (Biolegend) as previously reported (Herold et al. 2019). Quantitative RT-PCR was performed with SYBR Green mix (ABgene) on the instrument “Step One Realtime Cycler’’(ABgene) The RT-PCR program employed in this research is the following: 95°C for 15 min., 40x [95°C for 15 sec., 60°C for 20 sec. and 72°C for 15 sec.], 95°C for 15 sec. and 60°C for 60 sec. Relative expression was generally calculated with ΔΔCt relative quantification method. Melt curve was performed for all primers. Primers used for this publication are listed.

### Immunoblot and CO-Immunoprecipitation

Cells have been lysed in RIPA lysis buffer (20 mM Tris-HCl pH 7.5, 150 mM NaCl, 1mM Na^2^EDTA, 1mM EGTA, 1% NP-40 and 1% sodium deoxycholate), containing proteinase inhibitor (1/100) via sonication with Branson Sonifier 250 (duty cycle at 20% and output control set on level 2; 10 sonication / 1 minute cycles per sample). Protein concentration was quantified using Bradford assay as previously described (Prieto-Garcia et al., 2020a). 50μg protein was boiled in 5x Laemmli buffer (312.5mM Tris-HCl pH 6.8, 500 mM DTT, 0.0001% Bromphenol blue, 10% SDS and 50% Glycerol) for 5 min and separated on 10% Tris-gels in Running buffer (1.25M Tris base, 1.25M glycine and 1% SDS). After separation, protein was transferred to Polyvinylidene difluoride membranes (Immobilon-FL) in Transfer Buffer (25mM Tris base, 192mM glycine and 20% methanol). Membrane was exposed to blocking buffer (0.1% casein, 0.2xPBS and 0.1% Tween20) for 45 min at room temperature (RT). Then, membranes were incubated with listed primary Abs (1/1000 dilution in a buffer composed by 0.1% casein, 0.2x PBS and 0.1% Tween20) for 6h at room temperature (RT). Indicated secondary Abs (1/10000 dilution in a buffer composed by 0.1% casein, 0.2x PBS, 0.1% Tween20 and 0.01% SDS) were incubated for 1h at RT. Membranes were recorded in Odyssey® CLx Imaging System, and analysed using Image Studio software (Licor Sciences).

Immunoprecipitation was performed using 0.25 mg of Pierce™ Protein A/G Magnetic Beads (ThermoFisher), 1μg of the listed specific Ab and 500μg of protein lysate. For endogenous Co-Immunoprecipitations, beads were incubated with IgG (Sigma) as a control for specificity. Chromatin fractionation was performed adding 1% Triton X-100 to the lysis buffer as previously described (Parisis Nikos; Labome; 2013). TUBE assay was performed as previously indicated (Prieto-Garcia et al., 2020a), 100 μg/ml recombinant expressed GST-4x UIM-Ubiquilin fusion protein was added to the cells before protein extraction. Antibodies and dilutions used for this publication are listed.

### Immunohistochemistry and Immunofluorescence

For IF and IHC, primary antibodies were incubated ON at 4°C, followed by subsequent incubation with the secondary antibody for 1 hour at room temperature. After antibody exposure, slides were washed twice with PBS. Stained samples were mounted with Mowiol®40-88. IHC were recorded using Pannoramic DESK scanner and analyzed with Case Viewer software (3DHISTECH). For IF, tissue-samples/cells were counterstained with 5 μg/ml DAPI for 15 minutes after secondary antibody application. IF stained slides were recorded using a FSX100 microscopy system (Olympus). For antibodies, manufacturer’s manuals and instructions regarding concentration or buffer solutions were followed. TP53BP1 foci were analysed in 5 regions of interest (ROI, 10 cells per field) using ImageJ. For ɣ-H2AX, nuclear intensity was measured using ImageJ or the Operetta CLS High-Content Analysis System (Perkin Elmer). Number of cells or fields analysed were indicated

### Cell viability, Operetta system and IC_50_/GI_50_ calculation

For cell viability, cells were stained with 0,5% Crystal violet and analyzed using ImageJ software (staining intensity is between 0 to 255). Upon quantification of the staining intensity, values were normalized to control. Number of cells was quantified using Operetta High-Content Imaging System (PerkinElmer) (number of DAPI+ cells) or Invitrogen Countess II FL (number of cells after trypsinization) upon indicated treatments. For the Operetta High-Content Imaging System, cells were seeded in 384-well plates at equal density and exposed to indicated treatments. Then, cells were fixed using 4% PFA for 10 minutes and then, permealized using 0,5% Triton x100 in PBS for 5 minutes. Before quantification cells were stained with DAPI. Number of cells was determined counting the number of nucleus with the Harmony Software (Perkin Elmer). Loewe synergy as calculated using the Combenefit software as previously described (Di Veroli GY et al 2016). For the quantification, unhealthy cells with modified nuclear morphology were excluded. IC_50_ was calculated and visualized using the website: www.aatbio.com

### sgRNA and shRNA Design

sgRNAs were designed using the CRISPRtool (https://zlab.bio/guide-design-resources). shRNA sequences were designed using SPLASH-algorithm (http://splashrna.mskcc.org/) (Pelossof et al., 2017) or the RNAi Consortium/Broad Institute (www.broadinstitute.org/rnai-consortium/rnai-consortium-shrna-library).

### AAV and lentivirus production and purification

Viruses were synthetized in HEK293-T cells. For AAV production, cells were co-transfected with the plasmid of interest (10 μg), pHelper (15 μg) and pAAV-DJ (10 μg) using PEI (70 μg). AAV Virus isolation from transfected cells was performed as previously described (Prieto-Garcia et al. 2019). For Lentivirus production, HEK293 cells (70% confluence) were transfected with the plasmid of interest (15 μg), pPAX (10 μg) and pPMD2 (10 μg) using PEI (70 μg). After 96 H, the medium containing lentivirus was filtered (0.45 μM) and stored at −80°C.

### In Vivo Experiments and Histology

All *in vivo* experiments were approved by the Regierung Unterfranken and the ethics committee under the license numbers 2532-2-362, 2532-2-367, 2532-2-374 and 25322-1003. The mouse strains used for this publication are listed. All animals are housed in standard cages in pathogen-free facilities on a 12-h light/dark cycle with *ad libitum* access to food and water. FELASA2014 guidelines were followed for animal maintenance.

Adult mice were anesthetized with Isoflurane and intratracheally intubated with 50 μl AAV virus (3 × 10^7^ PFU) as previoulsy decribed (Prieto-Garcia et al. 2019). Viruses were quantified using Coomassie staining protocol(Kohlbrenner et al., 2012). Animals were sacrificed by cervical dislocation and lungs were fixed using 5% NBF. For IHC and H&E, slides were de-paraffinized and rehydrated following the protocol: 2x 5 min. Xylene, 2x 3 min. EtOH (100%), 2x 3 min. EtOH (95%), 2x 3 min. EtOH (70%), 3 min. EtOH (50%) and 3 min. H_2_O. For all staining variants, slides were mounted with 200 μl of Mowiol® 40-88 covered up by a glass coverslip. IHC slides were recorded using Pannoramic DESK scanner or using FSX100 microscopy system (Olympus) and analysed using Case Viewer software (3DHISTECH) and ImageJ. IF samples were recorded using FSX100 microscopy system (Olympus)

### RNA-sequencing

RNA sequencing was performed with Illumina NextSeq 500 as described previously (Buchel et al., 2017).RNA was isolated using ReliaPrep™ RNA Cell Miniprep System Promega kit, following the manufacturer’s instruction manual. mRNA was purified with NEBNext® Poly(A) mRNA Magnetic Isolation Module (NEB) and the library was generated using the NEBNext® UltraTM RNA Library Prep Kit for Illumina, following the manufacturer’s instructions).

### Sample preparation for mass spectrometry

The sample preparation was performed as described previously(Klann et al., 2020). Briefly, lysates were precipitated by methanol/chloroform and proteins resuspended in 8 M Urea/10 mM EPPS pH 8.2. Concentration of proteins was determined by Bradford assay and 100 μg of protein per samples was used for digestion. For digestion, the samples were diluted to 1 M Urea with 10mM EPPS pH 8.2 and incubated overnight with 1:50 LysC (Wako Chemicals) and 1:100 Sequencing grade trypsin (Promega). Digests were acidified using TFA and tryptic peptideswere purified by tC18 SepPak (50 mg, Waters). 125 μg peptides per sample were TMT labelled and the mixing was normalized after a single injection measurement by LC-MS/MS to equimolar ratios for each channel. 250 μg of pooled peptides were dried for offline High pH Reverse phase fractionation by HPLC.

### Offline high pH reverse phase fractionation

Peptides were fractionated using a Dionex Ultimate 3000 analytical HPLC. 250 μg of pooled and purified TMT-labeled samples were resuspended in 10 mM ammoniumbicarbonate (ABC), 5% ACN, and separated on a 250 mm long C18 column (X-Bridge, 4.6 mm ID, 3.5 μm particle size; Waters) using a multistep gradient from 100% Solvent A (5% ACN, 10 mM ABC in water) to 60% Solvent B (90% ACN, 10 mM ABC in water) over 70 min. Eluting peptides were collected every 45 s into a total of 96 fractions, which were cross-concatenated into 12 fractions and dried for further processing.

### LC-MS^3^ proteomics

All mass spectrometry data was acquired in centroid mode on an Orbitrap Fusion Lumos mass spectrometer hyphenated to an easy-nLC 1200 nano HPLC system using a nanoFlex ion source (ThermoFisher Scientific) applying a spray voltage of 2.6 kV with the transfer tube heated to 300°C and a funnel RF of 30%. Internal mass calibration was enabled (lock mass 445.12003 m/z). Peptides were separated on a self-made, 32 cm long, 75μm ID fused-silica column, packed in house with 1.9 μm C18 particles (ReproSil-Pur, Dr. Maisch) and heated to 50°C using an integrated column oven (Sonation). HPLC solvents consisted of 0.1% Formic acid in water (Buffer A) and 0.1% Formic acid, 80% acetonitrile in water (Buffer B).

For total proteome analysis, a synchronous precursor selection (SPS) multi-notch MS3 method was used in order to minimize ratio compression as previously described (McAlister et al., 2014). Individual peptide fractions were eluted by a non-linear gradient from 4 to 40% B over 210 minutes followed by a step-wise increase to 95% B in 6 minutes which was held for another 9 minutes. Full scan MS spectra (350-1400 m/z) were acquired with a resolution of 120,000 at m/z 200, maximum injection time of 50 ms and AGC target value of 4 x 10^5^. The most intense precursors with a charge state between 2 and 6 per full scan were selected for fragmentation within 3 s cycle time and isolated with a quadrupole isolation window of 0.4 Th. MS2 scans were performed in the Ion trap (Turbo) using a maximum injection time of 50ms, AGC target value of 1 x 10^4^ and fragmented using CID with a normalized collision energy (NCE) of 35%. SPS-MS3 scans for quantification were performed on the 10 most intense MS2 fragment ions with an isolation window of 1.2 Th (MS) and 2 m/z (MS2). Ions were fragmented using HCD with an NCE of 65% and analyzed in the Orbitrap with a resolution of 50,000 at m/z 200, scan range of 100-200 m/z, AGC target value of 1.5 x10^5^ and a maximum injection time of 150ms. Repeated sequencing of already acquired precursors was limited by setting a dynamic exclusion of 60 seconds and 7 ppm and advanced peak determination was deactivated.

## QUANTIFICATION AND STATISTICAL ANALYSIS

### RNA-sequencing analysis

Fastq files were generated using Illuminas base calling software GenerateFASTQ v1.1.0.64 and overall sequencing quality was analyzed using the FastQC script. Reads were aligned to the human genome (hg19) using Tophat v2.1.1 (Kim and Salzberg, 2011) and Bowtie2 v2.3.2(Langdon, 2015) and samples were normalised to the number of mapped reads in the smallest sample. For differential gene expression analysis, reads per gene (Ensembl gene database) were counted with the “summarizeOverlaps” function from the R package “GenomicAlignments” using the “union”-mode and non- or weakly expressed genes were removed (mean read count over all samples <1). Differentially expressed genes were called using edgeR(Robinson et al., 2010) and resulting p-values were corrected for multiple testing by false discovery rate (FDR) calculations. GSEA analyses were done with signal2Noise metric and 1000 permutations. Reactome analysis were performed with PANTHER(Mi et al., 2013) using the “Statistical overrepresentation test” tool with default settings. Genes were considered significantly downregulated for reactome analysis when: Log2FC>0.75 and FDR p-value<0.05.

### Proteomics analysis

Proteomics raw files were processed using proteome discoverer 2.2 (ThermoFisher). Spectra were recalibrated using the Homo sapiens SwissProt database (2018-11-21) and TMT as static modification at N-terminus and Lysines, together with Carbamidomethyl at cysteine residues. Spectra were searched against human database and common contaminants using Sequest HT with oxidation (M) as dynamic modification together with methionine-loss + acetylation and acetylation at the protein terminus. TMT6 (N-term, K) and carbamidomethyl were set as fixed modifications. Quantifications of spectra were rejected if average S/N values were below 5 across all channels and/or isolation interference exceeded 50%. Protein abundances were calculated by summing all peptide quantifications for each protein.

Reactome analysis were performed with PANTHER using the “Statistical overrepresentation test” tool with default settings. Proteins were considered significantly downregulated for reactome analysis when: FC<-0.5 and p-value<0.05. Heatmap visualization was performed using Morpheus (Broad Institute).

### Analysis of publicly available data

All publicly available data and software used for this publication are listed (Appendix Table S5). Oncoprints were generated using cBioportal (Cerami et al., 2012; Gao et al., 2013). Briefly, Oncoprints generates graphical representations of genomic alterations, somatic mutations, copy number alterations and mRNA expression changes. TCGA data was used for the different analysis. Data were obtained using UCSC Xena (https://doi.org/10.1101/326470). Data was download as log2(norm_count+1)

Box plots using TCGA and GTEx data were generated using the online tool BoxPlotR (Spitzer et al., 2014) and GEPIA(Tang et al., 2017). For BoxplotR, the data previously download from UCSC Xena was used to generate the graphics, p-values were calculated using two-tailed t-test. For Gepia. The differential analysis was based on: “TCGA tumors vs (TCGA normal)”, whereas the expression data were log2(TPM+1) transformed and the log2FC was defined as median(tumor) – median(normal). p-values were calculated with a one-way ANOVA comparing tumor with normal tissue. Tukey and Altman whiskers where used depending of the number of samples. Correlation analysis were calculated using using GEPIA’s software. The analysis was based on the expression of the following datasets: “TCGA tumors”, “TCGA normal”. p-values for correlation coefficents were calculated using two-tailed Student’s t-tests.

Heatmap Genomic signature comparing primary human lung tumor samples was performed using UCSC Xena (https://doi.org/10.1101/326470) based on the dataset “TCGA tumors”. Compared Gene Expression across different cell lines was perfomed using the online tool R2.

## Supporting information

STAR Methods

## DATA AND SOFTWARE AVAILABILITY

RNA-sequencing data is available at the Gene Expression Omnibus under the accession number GEO: GSE129982.

## Contact for reagent and resource sharing

Further information and requests for resources and reagents should be directed to and will be fulfilled by the Lead Contact, Markus E. Diefenbacher.

## Acknowledgements

We are grateful to the animal facility and Barbara Bauer at the Biocenter, University Würzburg. C.P.G. and O.H. are supported by the German Cancer Aid via grant 70112491, M.R. is funded by the DFG-GRK 2243 and IZKF B335. M.E.D. and M.R. are funded by the German Israeli Foundation grant 1431. T. F. is funded by the IZKF program Z2/CS-1.

## Author contributions

Conceptualization: C.P.G, M.E.D.; Methodology: C.P.G. (in vitro) and O.H. (*in vivo*), C.P.G. (Biochemistry), M.Re. (TUBE), C.S.V. (Operetta system) K.K. and Ch. Mü. (Mass Spec); CV.M. (ChIP); Formal analysis: C.P.G. (Bioinformatics), M.Ro. and M.E.D. (Pathology), K.K. and Ch. Mü. (Mass Spec), I.D.; Investigation: C.P.G., O.H., M.Re., T.F., M.Ro., M.E.D. Resources: R.K., M.Ro., M.E.D.; Writing-original draft: M.E.D.; Writing-review and editing: C.P.G., O.H., M.Re., M.Ro, K.K., I.D., M.E.D.; Supervision: M.E.D.; Funding acquisition: M.E.D.

## Conflict of Interest

The authors declare no potential conflicts of interest.

## Notes

### Competing Interest Statement

The authors have declared no competing interest.

