## Supplementary material for "Inhibition of USP28 overcomes Cisplatin-Resistance of Squamous Tumors by Suppression of the Fanconi Anemia Pathway": STAR Methods

**Material and Methods Table**

| **1^st^ Antibodies** | **Company** | **Identifier** | **RRID** |
| --- | --- | --- | --- |
| Polyclonal rabbit anti-USP28 | Sigma-Aldrich | HPA006778 | AB_1080520 |
| Polyclonal rabbit anti-USP28 | Sigma-Aldrich | HPA006779 | AB_1080517 |
| Rabbit monoclonal anti-p-ATM (ser1981) | Cell signalling | 13050 | AB_10835213 |
| Monoclonal mouse anti p-ATM | Santa Cruz | sc-47739 | AB_781524 |
| Monoclonal mouse anti-ACTIN/ B-ACTIN (C4) | Santa Cruz | sc-47778 | AB_626632 |
| Monoclonal mouse anti-VINCULIN (hVIN-1) | Sigma-Aldrich | V9131 | AB_477629 |
| Polyclonal rabbit anti-53BP1 | Santa Cruz | sc-22760 | AB_2256326 |
| Polyclonal rabbit anti p-ATR (ser428) | Cell signalling | 2853 | AB_2290281 |
| Monoclonal rabbit anti-p-H2a.x (ser139) | Cell signalling | 2577 | AB_2118010 |
| Monoclonal rabbit anti-p-H2a.x (ser139) | Millipore | 05-636 | AB_309864 |
| Monoclonal mouse anti-TUBULIN | Proteintech Europe | 66031-1-lg | AB_11042766 |
| Polyclonal rabbit anti-H3 | Abcam | ab18521 | AB_732917 |
| Polyclonal rabbit anti p-USP28 (ser67) | ThermoFisher | PA5-64727 | AB_2664483 |
| Polyclonal rabbit anti p-USP28 (ser714) | ThermoFisher | PA5-64728 | AB_2664484 |
| Polyclonal rabbit anti-P63 | Biolegend | 619001 | AB_2256361 |
| Monoclonal rabbit anti-p63 recombinant | Bimake | A5182 |  |
| Polyclonal rabbit anti-cleaved caspase 3 (Asp175) | Cell signalling | 9661 | AB_2341188 |
| Monoclonal mouse anti c-JUN | Santa Cruz | sc-74543 | AB_1121646 |
| Polyclonal rabbit anti-c-MYC (N-262) | Santa Cruz | sc-764 | AB_631276 |
| Monoclonal mouse anti FANCD2 | Abcam | ab108928 | AB_10862535 |
| Polyclonal rabbit anti P53BP1 | NOVUS | NB100-904 | AB_10002714 |
| Polyclonal rabbit anti P-P53 (ser15) | Cell signalling | 9284 | AB_331464 |
| Monoclonal rabbit anti p-(Ser/Thr) ATM + ATR Substrate | ThermoFischer | MA5-14872 | AB_10978748 |
| Monoclonal rabbit anti cleaved caspase 9 | Bimake | A5074 |  |
| Monoclonal mouse anti caspase 9 | Santa cruz | sc-73548 | AB_1120024 |
| Monoclonal rabbit anti RAD51 | Abcam | ab133534 | AB_2722613 |
| Monoclonal rabbit anti FANCI | Abcam | ab15344 | AB_443182 |

| **­2^nd^ Antibodies** | **Company** | **Identifier** | **RRID** |
| --- | --- | --- | --- |
| Donkey anti-Mouse IgG (H+L) Cross-Adsorbed Secondary Antibody, DyLight 680 | ThermoFisher | SA5-10170 | AB_2556750 |
| Donkey anti-Rabbit IgG (H+L) Cross-Adsorbed Secondary Antibody, DyLight 680 | ThermoFisher | A32802 | AB_2762836 |
| Donkey anti-Rabbit IgG (H+L) Cross-Adsorbed Secondary Antibody, DyLight 800 | ThermoFisher | SA5-10044 | AB_2556624 |
| Donkey anti-Mouse IgG (H+L) Cross-Adsorbed Secondary Antibody, DyLight 800 | ThermoFisher | SA5-10172 | AB_2556752 |
| Donkey anti-Goat IgG (H+L) Cross-Adsorbed Secondary Antibody, DyLight 800 | ThermoFisher | SA5-10092 | AB_2556672 |
| Donkey anti-Rabbit IgG (H+L) Highly Cross-Adsorbed Secondary Antibody, Alexa Fluor 488 | ThermoFisher | A21206 | AB_2535792 |
| Donkey anti-Mouse IgG (H+L) Highly Cross-Adsorbed Secondary Antibody, Alexa Fluor 488 | ThermoFisher | A21202 | AB_141607 |
| Donkey anti-Mouse IgG (H+L) Highly Cross-Adsorbed Secondary Antibody, Alexa Fluor 555 | ThermoFisher | A31570 | AB_2536180 |
| Donkey anti-Rabbit IgG (H+L) Highly Cross-Adsorbed Secondary Antibody, Alexa Fluor 555 | ThermoFisher | A31572 | AB_162543 |
| Donkey anti-Rabbit IgG (H+L) Highly Cross-Adsorbed Secondary Antibody, Alexa Fluor 647 | ThermoFisher | A-31573 | A  B_2536183 |

| **Bacterial Strains** | **Company** | **Identifier** |
| --- | --- | --- |
| **DH5α** F- endA1 glnV44 thi-1 recA1 relA1 gyrA96 deoR nupG Φ80dlacZΔM15 Δ(lacZYA-argF)U169, hsdR17(rK-mK+), λ– | ThermoFisher | 18263012 |
| **Chemicals and Commercial Assays** | **Company** | **Identifier** |
| Gibco™  Dulbecco’s Modified Eagle Medium (DMEM), high glucose | ThermoFisher | 11574486 |
| Gibco™ RPMI 1640 Medium | ThermoFisher | 21875158 |
| Gibco™ Trypsin-EDTA (0.5%), No Phenol Red | ThermoFisher | 15400054 |
| Fetal Bovine Serum (FCS) | Sigma-Aldrich | 12103C |
| Penicillin-Streptomycin | Sigma-Aldrich | P4333 |
| Gibco™ GlutaMAX™ Supplement | ThermoFisher | 35050061 |
| Polybrene | Sigma-Aldrich | TR-1003 |
| Polyethylenimine, Linear, MW 25000, Transfection Grade (PEI 25K) | Polysciences | 23966-1 |
| Dimethyl sulfoxide (DMSO) | Sigma-Aldrich | D8418 |
| Ethanol (Etoh) | Carl Roth | 5054.6 |
| Gibco™ Phosphate-buffered saline (PBS) | ThermoFisher | 10010031 |
| Nuclease-free water | Merck | 3098 |
| Cycloheximide | Sigma-Aldrich | 1810 |
| Doxycycline hyclate | Sigma-Aldrich | D9891 |
| Tandem ubiquitin binding entity (TUBE) | Homemade |  |
| PR-619 | Selleckchem | S7130 |
| AZ1 | Probechem | PC-60023 |
| Propidium Iodide (PI) | Sigma-Aldrich | 11348639001 |
| Protease Inhibitor Cocktail | Roche | 4693159001 |
| Pierce™ Protein A/G Magnetic Beads | ThermoFisher | 88802 |
| RIPA Lysis Buffer | Homemade |  |
| 4′,6-diamidino-2-phenylindole (DAPI) | ThermoFisher | D1306 |
| Hoechst | ThermoFisher | 62249 |
| Coomassie Brilliant Blue R-250 Dye | ThermoFisher | 20278 |
| Phosphate-buffered saline (PBS) | Homemade |  |
| 2-Propanol/ Isopropanol | ROTH | AE73.2 |
| Adenosintriphosphat (ATP) | Jena Bioscience | NU-1010-10G |
| Agarose | ROTH | 3810.4 |
| Ampicillin (Amp) | ROTH | HP62.2 |
| Bovine serum albumine (BSA) | Merck Millipore | 810683 |
| Mowiol® 40-88 | Sigma-Aldrich | 324590 |
| Dithiothreitol (DTT) | Sigma-Aldrich | D9779 |
| Eosin | Sigma | E4009 |
| Hematoxylin | Sigma | H3136 |
| Polyvinylidene difluoride membranes (PVDF) Immobilon Transfer Membrane | Merck | IPFL00010 |
| N,N,N',N'-tetramethylenethylendiamine (TEMED) | ROTH | 2367.3 |
| Natrium chloride (NaCl) | AppliChem | A2942,1000 |
| Neutrally buffered formalin (NBF) | Thermo Fisher | 5700TS |
| Tris-HCl | ROTH | 9090.5 |
| TritonX100 | ROTH | 3051.3 |
| Xylene | Sigma | 534056 |
| β-Mercaptoethanol | ROTH | 4227.1 |
| Methanol (MeOH) | ROTH | 0082.3 |
| Transfer Buffer (Westerm Blot) | Homemade |  |
| Blocking Buffer (Western Blot) | Homemade |  |
| Wash Buffer (western Blot( | Homemade |  |
| SDS Loading Buffer/ lämmli buffer (5X) | Homemade |  |
| SDS Running Buffer (Western Blot) | Homemade |  |
| Primary antibody buffer (Western Blot) | Homemade |  |
| Secondary antibody buffer (western Blot) | Homemade |  |
| peq GOLD Trifast | VWR (Peqlab brand) | 30-2010 |
| M-MLV reverse transcriptase | Promega | M1701 |
| M-MLV RT 5X Buffer | Promega | M531A |
| Deoxynucleotidetriphosphates (dNTPs) Mix | Promega | U151A |
| Random Hexamer Primer | ThermoFisher | SO142 |
| RiboLock RNase Inhibitor | ThermoFisher | EO0381 |
| SYBR™ Green PCR Master Mix | ThermoFisher | 4309155 |
| ReliaPrep™ RNA Cell Miniprep System Protocol | Promega | TM370 |
| NEBNext® Ultra™RNA Library Prep Kit for Illumina | New England Biolabs (NEB) | NEB #E7530S |
| NEBNext® Multiplex Oligos for Illumina® (Dual Index Primers Set 1) | New England Biolabs (NEB) | NEB #E7600S |
| NEBNext® Poly(A) mRNA Magnetic Isolation Module | New England Biolabs (NEB) | NEB #E7490S |
| N,N-Dimethylformamid (DMF) | Sigma | 61747 |
| Cisplatin | Selleckhem | S1166 |
| 5-Fluorouracil | Selleckhem | S1209 |
| Oxaliplatin | Selleckhem | S1224 |
| VE-821 | Selleckhem | S8007 |
| KU‑55933 | Selleckhem | S1092 |
| **Cell lines**  **Cell** | **Company** | **Identifier** |
| Human: HEK 293T | ATCC | ATCC® CRL-11268™ |
| Human: A-431 | ATCC | ATCC® CRL-1555 |
| Human: LUDLU-1 | ECACC | 92012463 |
| Human: H1299 | ATCC | ATCC® CRL-5803 |
| Human: HELA | ATCC | ATCC® CCL-2 |
| Human: SiHa | ATCC | ATCC® HTB-35 |
| Human: Ca Ski | ATCC | ATCC® CRL-1550 |
| Human: BEAS2-B | ATCC | ATCC® CRL-9609 |
| Human: CALU1 | ATCC | ATCC® HTB-54 |
| Human: Detroit 562 | ATCC | ATCC® CCL-138 |
| Human: SK-MES1 | ATCC | ATCC® HTB-58 |
| Human: H23 | ATCC | ATCC® CRL-5800 |
| Mouse: KP ADC | Primary tumors |  |
| Mouse: KPL SCC | Primary tumors |  |
| **Experimental Models: Organisms/Strains Cell** | **Company** | **Identifier** |
| B6(C)-Gt(ROSA)26Sor^em1.1(CAG-cas9*,-EGFP)Rsky^/J | The Jackson laboratory | Stock No: 028555 |
| B6.129-Kras^tm4Tyj^ Trp53^tm1Brn^/J | The Jackson laboratory | Stock No: 032435 |
| C57BL/6J | The Jackson Laboratory | Stock No: 000664 |
| **Oligonucleotides** | **Sequence** | **Company** |
| hUsp28 FW | ACTCAGACTATTGAACAGATGTACTGC | Sigma |
| hUsp28 RV | CTGCATGCAAGCGATAAGG | Sigma |
| hFANCI FW | TTTGCCATCAAATTGGACTATG | Sigma |
| hFANCI RV | TTGGAATCTCCTTGCTGTCC | Sigma |
| hB-Actin FW | GCTACGAGCTGCCTGACG | Sigma |
| hB-Actin RV | GGCTGGAAGAGTGCCTCA | Sigma |
| hFANCD2 FW | CCCAGAACTGATCAACTCTCCT | Sigma |
| hFANCD2 RV | CCATCATCACACGGAAGAAA | Sigma |
| hTp63 FW | GGAAAACAATGCCCAGACTC | Sigma |
| hTp63 RV | GTGGAATACGTCCAGGTGGC | Sigma |
| hTp63-2 FW | GAAAGCTGTTCCTTGGTCCTAGT | Sigma |
| hTP63-2 RV | GGTTTATTCAAACCCTCAGCA | Sigma |
| hRAD51C FW | TGGATTTGGTGAGTTTCCCGC | Sigma |
| hRAD51C RV | TCTTTGCTAAGCTCGGAGGG | Sigma |
| mUSP28 FW | ATGACAACTTGCCCCACTTC | Sigma |
| mUSP28 RV | AGTTCCACAGACAGGGCTTC | Sigma |
| mB-Actin FW | AGTGTGACGTTGACATCCGT | Sigma |
| mB-Actin RV | TGCTAGGAGCCAGAGCAGTA | Sigma |
| human shRNA ∆Np63 #1 for | TGAATGAACAGACGTCCAATTTCTCGAGAAATTGGACGTCTGTTCATTCTTTTTC | Sigma |
| human shRNA ∆Np63 #1 rev | TCGAGAAAAAGAATGAACAGACGTCCAATTTCTCGAGAAATTGGACGTCTGTTCATTCA | Sigma |
| human shRNA ∆Np63 #2 for | TCGAGTGGAATGATTTCAACTTCTCGAGAAGTTGAAATCATTCCACTCGTTTTTC | Sigma |
| human shRNA ∆Np63 #2 rev | TCGAGAAAAACGAGTGGAATGATTTCAACTTCTCGAGAAGTTGAAATCATTCCACTCGA | Sigma |
| human shRNA USP28 #1 for | CCGGCAAGGAGCTTATTCGAAATCTCGAGATTTCGAATAAGCTCCTTGTTTTTG | Sigma |
| human shRNA USP28 #1 rev | AATTCAAAAACAAGGAGCTTATTCGAAATCTCGAGATTTCGAATAAGCTCCTTG | Sigma |
| human shRNA USP28 #2 for | CCGGGACTGAAGATCATCCATTACTCGAGTAATGGATGATCTTCAGTCTTTTTG | Sigma |
| human shRNA USP28 #2 rev | AATTCAAAAAGACTGAAGATCATCCATTACTCGAGTAATGGATGATCTTCAGTC | Sigma |
| murine shRNA USP28 #1 for | TGCTGTTGACAGTGAGCGAGGATGTGAATTTGTATAAAAATAGTGAAGCCACAGATGTATTTTTATACAAATTCACATCCCTGCCTACTGCCTCGGA | Sigma |
| murine shRNA USP28 #2 for | TGCTGTTGACAGTGAGCGATCTGTTTATACTTTAGATAAATAGTGAAGCCACAGATGTATTTATCTAAAGTATAAACAGACTGCCTACTGCCTCGGA | Sigma |
| sgRNA murine Stk11/Lkb1 for | CACCGCGAGACCTTATGCCGCAGGG | Sigma |
| sgRNA murine Stk11/Lkb1 rev | AAACCCCTGCGGCATAAGGTCTCGc | Sigma |
| sgRNA murine Usp28 #1 for | CACCGGGGAGCCTTCCGATCATCCG | Sigma |
| sgRNA murine Usp28 #1 rev | AAACCGGATGATCGGAAGGCTCCCc | Sigma |
| sgRNA murine Usp28 #2 for | CACCGCGGATCGTTCCGTGAAGTAT | Sigma |
| sgRNA murine Usp28 #2 rev | AAACATACTTCACGGAACGATCCGc | Sigma |
| hUSP28S67A- for | ATGAGAGAGTTAAGGAGCCCGCTCAAGACACTGTTGCTACAGA | Sigma |
| hUSP28S67A-rev | TCTGTAGCAACAGTGTCTTGAGCGGGCTCCTTAACTCTCTCAT | Sigma |
| hUSP28S714A-for | AGTCCTCCACCAACTCCTCAGCACAGGACTACTCTACATCACA | Sigma |
| hUSP28S714A-rev | TGTGATGTAGAGTAGTCCTGTGCTGAGGAGTTGGTGGAGGACT | Sigma |
| hUsp28 S67A gblock rev BamHI | AAGCTTGGATCCTTACGTGCTTAGAATTGTGCCTG | Sigma |
| hUsp28 S714A gblock rev XbaI | AAGCTTTCTAGATCACATTCTAATGCCACAATTC | Sigma |
| ChIP FANCD2 distal For | GCTGTCTGGCAAGTTAGG A TGG | Sigma |
| ChIP FANCD2 distal Rev | CAAGCTGTAAGGCATTTCC CCG | Sigma |
| ChIP FANCI For | GGCGGATCTTGTTGTTACGG | Sigma |
| ChIP FANCI Rev | CCTCCGCCACAAACTTCCAA | Sigma |
| ChIP Rad51C For | TTTGGGGAATCAAAACGGAATGG | Sigma |
| ChIP Rad51C Rev | AGGCTCACCTGCTAACCCC | Sigma |
| sgRNA murine Kras #1 for | CACCGACTGAGTATAAACTTGTGG | Sigma |
| sgRNA murine Kras #1 rev | AAACCCACAAGTTTATACTCAGTC | Sigma |
| sgRNA murine Trp53 #1 for | CACCGATGGTGGTATACTCAGAGC | Sigma |
| sgRNA murine Trp53 #1 rev | AAACGCTCTGAGTATACCACCATC | Sigma |
| KrasG12D repair template for | TTTTGTGTAAGCTTTGGTAACTCCATGTATTTTTATTAAGTGTT | Sigma |
| KrasG12D repair template rev | GAGCTTATCGATACCGTCGACACACCCAGTTTAAAGCCTTGGAA | Sigma |
| **Recombinant DNA** | **Company/Source** | **Identifier** |
| pLKO.DEST.EGFP | pLKO.DEST.EGFP was a gift from Ming-Sound Tsao (Addgene plasmid # 32684 ; http://n2t.net/addgene:32684 ; RRID:Addgene_32684) | Addgene plasmid # 32684 |
| pLKO.1 puro | pLKO.1 puro was a gift from Bob Weinberg (Addgene plasmid # 8453 ; http://n2t.net/addgene:8453 ; RRID:Addgene_8453) | Addgene plasmid # 8453 |
| pINDUCER20 | pInducer20 was a gift from Stephen Elledge (Addgene plasmid # 44012 ; http://n2t.net/addgene:44012 ; RRID:Addgene_44012) | Addgene plasmid # 44012 |
| deltaNp63alpha-FLAG | deltaNp63alpha-FLAG was a gift from David Sidransky (Addgene plasmid # 26979 ; http://n2t.net/addgene:26979 ; RRID:Addgene_26979) | Addgene plasmid # 26979 |
| pLKO-eGFP-shdeltaNp63-1 (GFP) | This publication | N/A |
| pLKO-eGFP-shdeltaNp63-2 (GFP) | This publication | N/A |
| pDZ Flag USP28 | pDZ Flag USP28 (Addgene plasmid # 15665 ; http://n2t.net/addgene:15665 ; RRID:Addgene_15665) | Addgene plasmid # 15665 |
| pINDUCER20 -mouse Usp28 WT (Neomycin) | This publication | N/A |
| pLKO shUSP_28_1 (Puromycin) | This publication | N/A |
| pLKO shUSP_28_2 (Puromycin) | This publication | N/A |
| pcDNA3-HA-USP28 | pcDNA3-HA-USP28 was a gift from Nikita Popov | N/A |
| pGEPIR 20-human-sh-Usp28 (Neomycin) | pINDUCER 20-human-sh- Usp28 was a gift from Carina Maier (Almut Schulze group) | N/A |
| pLKO USP28 S67A | This publication | N/A |
| pLKO USP28 S714A | This publication | N/A |
| pLKO USP28 S61A + S714A | This publication | N/A |
| psPAX2 | psPAX2 was a gift from Didier Trono (Addgene plasmid # 12260 ; http://n2t.net/addgene:12260 ; RRID:Addgene_12260) | Addgene plasmid # 12260 |
| pMD2G | pMD2.G was a gift from Didier Trono (Addgene plasmid # 12259 ; http://n2t.net/addgene:12259 ; RRID:Addgene_12259) | Addgene plasmid # 12259 |
| pHelper | Cell Biolabs, INC. | VPK-400-DJ |
| pAAV2/8 | AAV2/8 was a gift from James M. Wilson (Addgene plasmid # 112864 ; http://n2t.net/addgene:112864 ; RRID:Addgene_112864) | Addgene plasmid # 112864 |
| pAAV-DJ Vector | Cell Biolabs, INC. | VPK-420-DJ |
| pLKO-eGFP-mshUSP_28_1 (GFP) | This Paper | N/A |
| pLKO--eGFP- mshUSP_28_2 (GFP) | This Paper | N/A |
| AAV:ITR- U6-sgRNA(p53)- pEFS-2A-mCherry-shortPA- ITR | This publication | N/A |
| AAV:ITR-U6-sgRNA(Kras)-pEFS-2A-mCherry-shortPA-KrasG12D_HDRdonor-ITR | This publication | N/A |
| AAV:ITR-U6-sgRNA(Kras)-U6-sgRNA(p53)-pEFS-2A-mCherry-shortPA-KrasG12D_HDRdonor-ITR | This publication | N/A |
| AAV:ITR-U6-sgRNA(Kras)-U6-sgRNA(p53)-U6-sgRNA(Lkb1)- pEFS-2A-mCherry-shortPA-KrasG12D_HDRdonor-ITR | This publication | N/A |
| AAV:ITR-U6-sgRNA(Kras)-U6-sgRNA(p53)-U6-sgRNA(Lkb1)-U6-sgRNA(Usp28^1^)-U6-sgRNA(Usp28^2^)-pEFS-2A-mCherry-shortPA-KrasG12D_HDRdonor-ITR | This publication | N/A |
| AAV:ITR-U6-sgRNA(Kras)-U6-sgRNA(p53)-U6-sgRNA(Lkb1)-pEFS-Rluc-2A-Cre-shortPA-KrasG12D_HDRdonor-ITR (AAV-KPL) | AAV:ITR-U6-sgRNA(Kras)-U6-sgRNA(p53)-U6-sgRNA(Lkb1)-pEFS-Rluc-2A-Cre-shortPA-KrasG12D_HDRdonor-ITR (AAV-KPL) was a gift from Feng Zhang (Addgene plasmid # 60224 ; http://n2t.net/addgene:60224 ; RRID:Addgene_60224) | Addgene plasmid # 60224 |
| **Software and Algorithm** | **Company/Source** |  |
| cBioportal | https://www.cbioportal.org |  |
| GEPIA and GEPIA2 | http://gepia.cancer-pku.cn |  |
| KM-plotter | http://kmplot.com/analysis/ |  |
| Operetta Imaging | Perkin Elmar |  |
| BoxPlotR | http://shiny.chemgrid.org/boxplotr/ |  |
| Excel | Microsoft |  |
| Affinity Desgner | https://affinity.serif.com/es/designer/ |  |
| Image Studio | Licor |  |
| Panther Classification system | http://pantherdb.org |  |
| AATBIO IC50 calculator | https://www.aatbio.com/tools/ic50-calculator |  |
| GraphPad Software | GraphPad Software, Inc. |  |
| Affinity Designer | Serif Europe |  |
| ImageJ | National Insistute of Health |  |
| Primerx | http://www.bioinformatics.org/primerx/cgi-bin/DNA_1.cgi |  |
| ROC Plotter | http://www.rocplot.org/ |  |
| Pannoramic Case Viewer | 3dHistech |  |
| R2: Genomics Analysis and Visualization Platform | http://r2.amc.nl |  |
| UCSC Xena | https://ucsc-xena.gitbook.io/project/ |  |
| GenerateFastq v1.1.0.64 | http://emea.support.illumina.com/downloads/local-run-manager-generate-fastq-module.html |  |
| FastQC | http://www.bioinformatics.babraham.ac.uk/projects/fastqc/ |  |
| Bowtie2 v2.3.4.1 | http://bowtie-bio.sourceforge.net/index.shtml |  |
| TopHat v.2.1.1 | <https://ccb.jhu.edu/software/tophat/index.shtml> |  |
| Samtools v1.3 | http://samtools.sourceforge.net |  |
| R | https://www.r-project.org |  |
| EdgeR | <https://bioconductor.org/packages/release/bioc/html/edgeR.html> |  |
| GenomicAlignments | <https://bioconductor.org/packages/release/bioc/html/GenomicAlignments.html> |  |
| GSEA v2.2 | <http://software.broadinstitute.org/gsea/downloads.jsp> |  |
| COMBENEFIT | <https://www.cruk.cam.ac.uk/research-groups/jodrell-group/combenefit> |  |
| EMBL | <https://www.embl.de/> |  |
| SPLASHRNA | <http://splashrna.mskcc.org/> |  |
| **Instrument** | **Company/Source** |  |
| Odyssey® CLx Imaging System | Licor |  |
| iBright™ FL1000 Imaging System | Invitrogen |  |
| BD FACSCanto II Cell Analyzer | BD Biosciences |  |
| StepOnePlus Real-Time PCR System | ThermoFisher |  |
| Invitrogen Countess II FL Automated Cell Counter | ThermoFisher |  |
| Pannoramic DESK scanner | 3DHISTECH |  |
| FSX100 microscopy | Olympus Life Science |  |
| Operetta screening and imaging system | Perkin Elmer |  |
| Fragment Analyzer | Agilent formerly Advanced Analytical |  |
| Axiocam 503 mono + Zeiss axio microscope | Zeiss |  |
| Branson Sonifier 250 | Branson |  |
| Hyrax M55 Rotary Microtome | Leica |  |
| PCR cycler: SimpliAmp thermo cycler | Life technologies |  |
| Orbitrap Fusion Lumos | ThermoFisher |  |
| Hyrax M55 Rotary Microtome | Leica |  |
| PCR cycler: SimpliAmp thermo cycler | Life technologies |  |
