## Supplementary material for "Inhibition of USP28 overcomes Cisplatin-Resistance of Squamous Tumors by Suppression of the Fanconi Anemia Pathway": Suppl. Figure 1-7

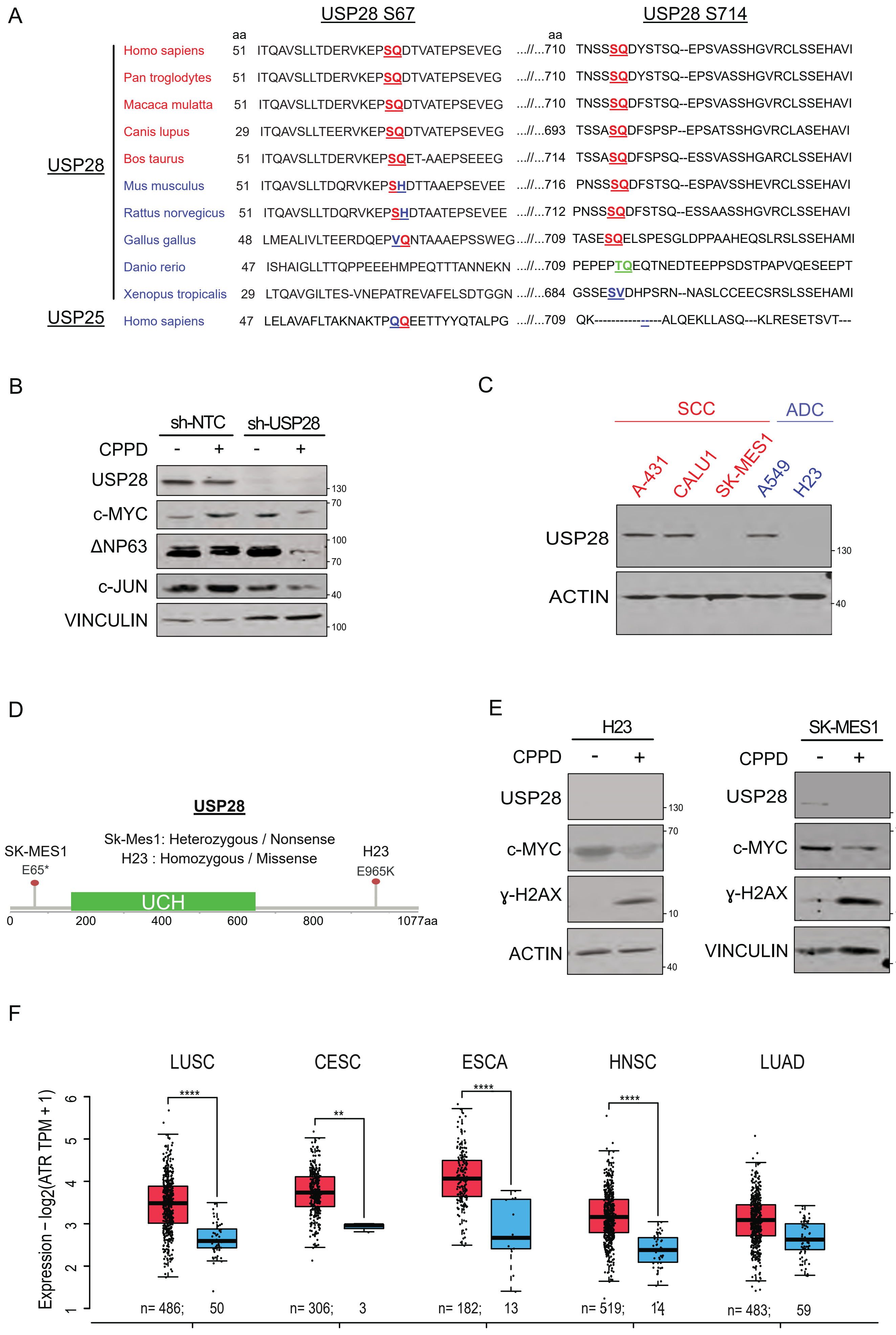

A
USP28 S67
USP28 S714
aa
aa
USP28
Homo sapiens 51 ITQAVSLLTDERVKEPSQDTVATEPSEVEG ...//...710 TNSSSQDYSTSQ--EPSVASSHGVRCLSSEHAVI
Pan troglodytes 51 ITQAVSLLTDERVKEPSQDTVATEPSEVEG ...//...710 TNSSSQDYSTSQ--EPSVASSHGVRCLSSEHAVI
Macaca mulatta 51 ITQAVSLLTDERVKEPSQDTVATEPSEVEG ...//...710 TNSSSQDFSTSQ--EPSVASSHGVRCLSSEHAVI
Canis lupus 29 ITQAVSLLTEERVKEPSQDTVATEPSEVEG ...//...693 TSSASQDFSPSP--EPSATSSHGVRCLASEHAVI
Bos taurus 51 ITQAVSLLTDERVKEPSQET-AAEPSEEEG ...//...714 TSSASQDFSPSQ--ESSVASSHGARCLSSEHAVI
Mus musculus 51 ITQAVSLLTDQRVKEPSHDTTAAEPSEVEE ...//...716 PNSSSQDFSTSQ--ESPAVSSHEVRCLSSEHAVI
Rattus norvegicus 51 ITQAVSLLTDQRVKEPSHDTAATEPSEVEE ...//...712 PNSSSQDFSTSQ--ESSAASSHGVRCLSSEHAVI
Gallus gallus 48 LMEALIVLTEERDQEPVQNTAAAEPSSWEG...//...709 TASESQELSPESGLDPPAAHEQSLRSLSSEHAMI
Danio rerio 47 ISHAIGLLTTQPPEEEHMPEQTTTANNEKN ...//...709 PEPEPTQEQTNEDTEEPPSDSTPAPVQESEEPT
Xenopus tropicalis 29 LTQAVGILTES-VNEPATREVAFELSDTGGN ...//...684 GSSESVDHPSRN--NASLCCEECSRSLSSEHAMI
USP25
Homo sapiens 47 LELAVAFLTAKNAKTPQQEETTYYQTALPG ...//...709 QK---------------ALQEKLLASQ---KLRESETSVT---
B
sh-NTC
sh-USP28
CPPD - + - +
USP28
c-MYC
ΔNP63
c-JUN
VINCULIN
130
70
100
70
40
100
C
SCC
ADC
A-431
CALU1
SK-MES1
A549
H23
USP28
ACTIN
130
40
D
USP28
Sk-Mes1: Heterozygous / Nonsense
H23 : Homozygous / Missense
SK-MES1
E65*
H23
E965K
UCH
0
200
400
600
800
1077aa
E
H23
SK-MES1
CPPD - +
CPPD - +
USP28
c-MYC
ɣ-H2AX
ACTIN
VINCULIN
130
70
10
40
100
F
LUSC
CESC
ESCA
HNSC
LUAD
Expression – log2(ATR TPM + 1)
1
2
3
4
5
6
****
**
****
****
n= 486; 50
n= 306; 3
n= 182; 13
n= 519; 14
n= 483; 59


Figure S1

***Figure S1 USP28 is recruited to DNA damage sites and phosphorylated by ATR, not ATM, upon Cisplatin treatment***

A) Sequence alignment within USP28 at serine 67 and serine 714 depicting inter-species conservation. Red=Conserved ATM/ATR phospho-motif respect to human; Blue=Non-conserved ATM/ATR phospho-motif respect to human; Green =SQ ATM/ATR phospho-motif was replaced for TQ ATM/ATR phospho-motif.

B) Immunoblotting against endogenous USP28, c-MYC, ΔNp63 and c-JUN in either control or stable shRNA-expressing USP28 knock down A431 cells, treated with either DMF or 5μM CPPD for 24 hours. ACTIN serves as loading control. n=3.

C) Immunoblotting against endogenous USP28 in SCC (A431, CALU1 and SK-MES1) and ADC (A549 and H23) cell lines. ACTIN serve as loading control. n=3.

D) Schematic representation of USP28 mutations presented in H23 and SK-MES1 lung cancer cell lines.

E) Immunoblotting against endogenous USP28, c-MYC and phospho-H2AX in the USP28 mutant cell line H23, treated with either DMF or 5μM CPPD for 24 hours. ACTIN serves as loading control. n=3.

F) ATR gene expression in tumour (red) and normal (blue) samples for lung SCC (LUSC), cervix SCC (CESC), head and neck SCC (HNSC) and lung ADC (LUAD) patient samples. ** $p<0.01$; **** $p<0.0001$. Data obtained from GEPIA2 online software.

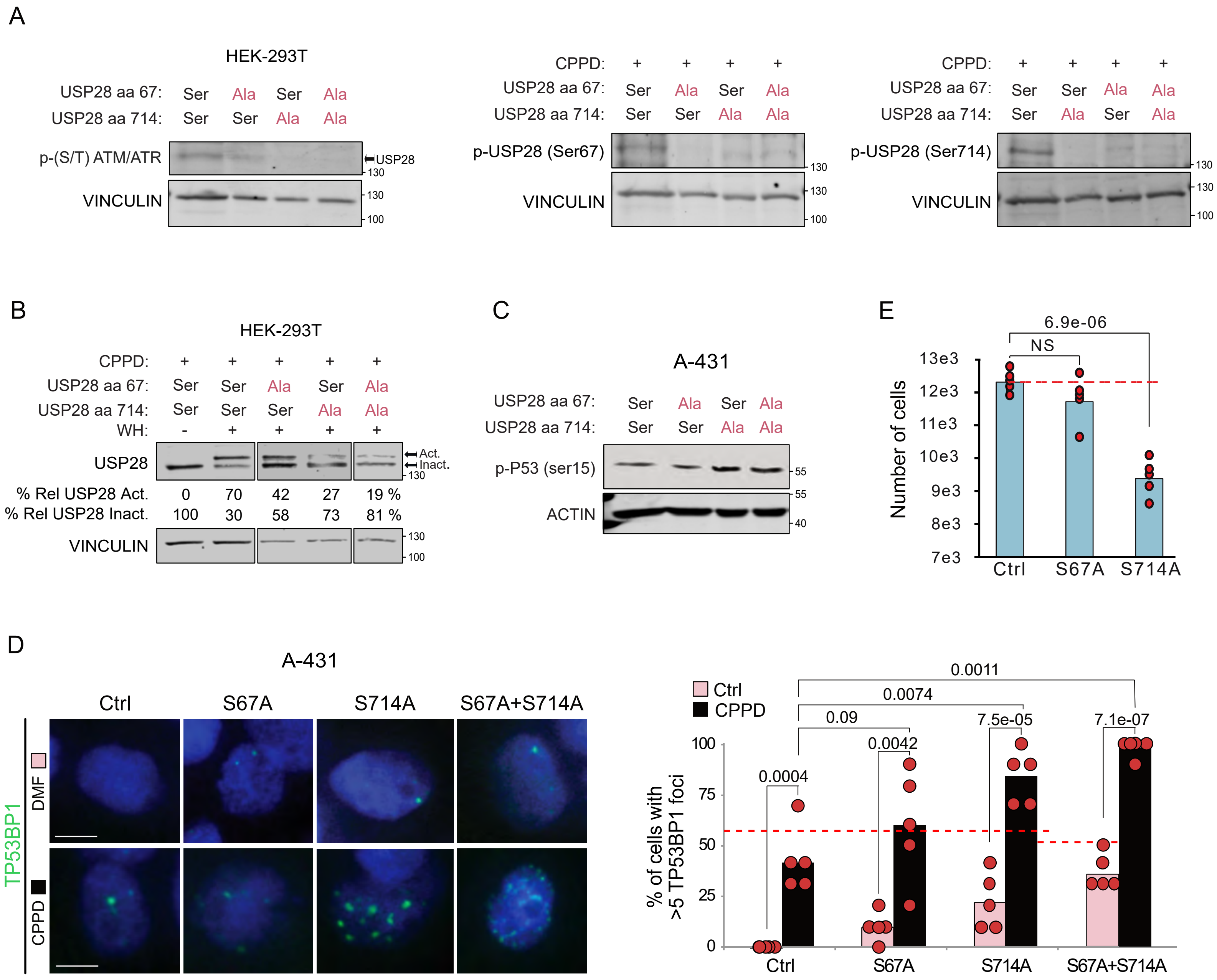

A
HEK-293T
USP28 aa 67: Ser Ala Ser Ala
USP28 aa 714: Ser Ser Ala Ala
p-(S/T) ATM/ATR
USP28
VINCULIN
CPPD: + + + +
USP28 aa 67: Ser Ala Ser Ala
USP28 aa 714: Ser Ser Ala Ala
p-USP28 (Ser67)
VINCULIN
CPPD: + + + +
USP28 aa 67: Ser Ser Ala Ala
USP28 aa 714: Ser Ala Ser Ala
p-USP28 (Ser714)
VINCULIN
B
HEK-293T
CPPD: + + + + +
USP28 aa 67: Ser Ser Ala Ser Ala
USP28 aa 714: Ser Ser Ser Ala Ala
WH: - + + + +
USP28
Act.
Inact.
% Rel USP28 Act. 0 70 42 27 19 %
% Rel USP28 Inact. 100 30 58 73 81 %
VINCULIN
C
A-431
USP28 aa 67: Ser Ala Ser Ala
USP28 aa 714: Ser Ser Ala Ala
p-P53 (ser15)
ACTIN
E
6.9e-06
NS
Number of cells
13e3
12e3
11e3
10e3
9e3
8e3
7e3
Ctrl
S67A
S714A
D
A-431
Ctrl
S67A
S714A
S67A+S714A
TP53BP1
DMF
CPPD
Ctrl
CPPD
0.0011
0.0074
0.09
7.5e-05
7.1e-07
0.0042
0.0004
% of cells with >5 TP53BP1 foci
100
75
50
25
0
Ctrl
S67A
S714A
S67A+S714A


Figure S2

***Figure S2 Phosphorylation of USP28 upon Cisplatin exposure is required to repair DNA damage in SCC***

A) Immunoblotting against ATR/ATM SQ/TQ motif, phosphorylated USP28 at serine 67 and serine 714, in control, S67A, S714A and S67A+S714A mutant HEK-293T cells treated with 5μM CPPD for 6 hours. VINCULIN serves as loading control. The arrow indicates the molecular weight of USP28. n=3

B) Ubiquitin suicide probe (warhead) assay, followed by immunoblotting against USP28 in control, S67A, S714A and S67A+S714A mutant HEK-293T cells exposed to 5 μM CPPD for 6 hours. 'Act.' arrow indicates active USP28. 'Inact.' arrow indicates inactive USP28. VINCULIN serves as loading control. n=3.

C) Immunoblotting against phosho-P53 at serine 15 in control, S67A, S714A and S67A+S714A mutant A431 cells treated with 5μM CPPD for 6 hours. ACTIN serves as loading control. n=3

D) Immunofluorescence against endogenous TP53BP1 in control, S67A, S714A and S67A+S714A mutant A431 cells, treated with either DMF or 5μM CPPD for 6 hours. DAPI served as nuclear marker. n=3. Percentage of cells with more than 5 P53BP1 foci was calculated measuring 10 cells per field (n=5). Scale bar= 10μm. T-Test was used to calculate the p-value.

E) Number of cells in control S67A and S714A mutant A431 cells, treated with DMF (blue) for 48 hours. Number of cells were calculated measuring 15 fields per well (n=5). P-values were calculated using two-tailed T-test statistical analysis.

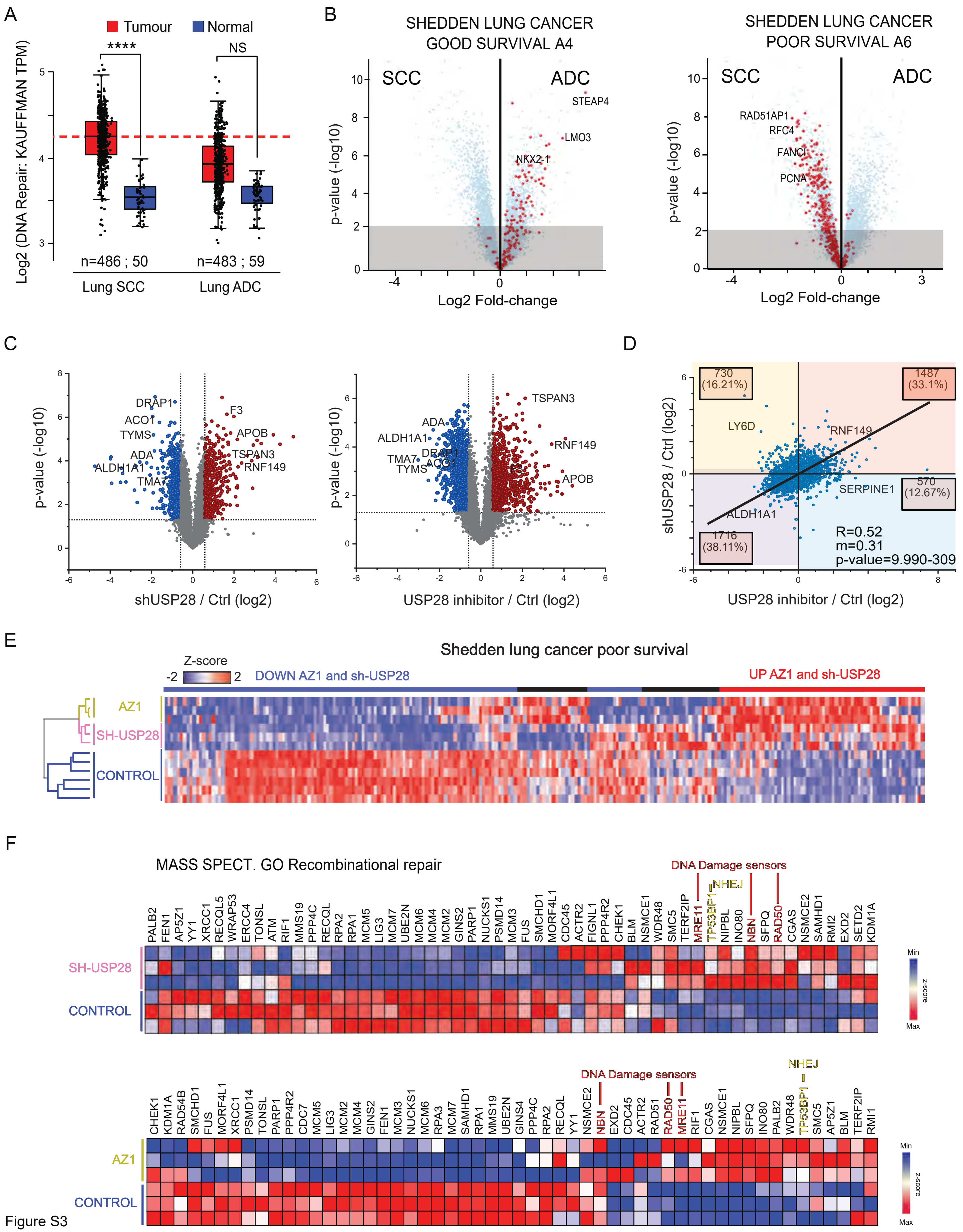

A
Tumour
Normal
****
NS
Log2 (DNA Repair: KAUFFMAN TPM)
n=486 ; 50
n=483 ; 59
Lung SCC
Lung ADC
B
SHEDDEN LUNG CANCER
GOOD SURVIVAL A4
SCC
ADC
STEAP4
LMO3
NKX2-1
p-value (-log10)
Log2 Fold-change
SHEDDEN LUNG CANCER
POOR SURVIVAL A6
RAD51AP1
RFC4
FANCI
PCNA
C
DRAP1
ACO1
TYMS
ADA
ALDH1A1
TMA7
F3
APOB
TSPAN3
RNF149
shUSP28 / Ctrl (log2)
USP28 inhibitor / Ctrl (log2)
D
730
(16.21%)
1487
(33.1%)
LY6D
570
(12.67%)
SERPINE1
1716
(38.11%)
R=0.52
m=0.31
p-value=9.990-309
shUSP28 / Ctrl (log2)
E
Shedden lung cancer poor survival
Z-score
DOWN AZ1 and sh-USP28
UP AZ1 and sh-USP28
AZ1
SH-USP28
CONTROL
F
MASS SPECT. GO Recombinational repair
DNA Damage sensors
NHEJ
PALB2 FEN1 AP5Z1 YY1 XRCC1 RECQL5 WRAP53 ERCC4 TONSL ATM RIF1 MMS19 PPP4C RECQL RPA2 RPA1 MCM5 LIG3 MCM7 UBE2N MCM6 MCM4 MCM2 GINS2 PARP1 NUCKS1 PSMD14 MCM3 FUS SMCHD1 MORF4L1 CDC45 ACTR2 FIGNL1 PPP4R2 CHEK1 BLM NSMCE1 WDR48 SMC5 TERF2IP MRE11 TP53BP1 NIPBL INO80 NBN SFPQ RAD50 CGAS NSMCE2 SAMHD1 RMI2 EXD2 SETD2 KDM1A
Min
Max
z-score
CHEK1 KDM1A RAD54B SMCHD1 FUS MORF4L1 XRCC1 PSMD14 TONSL PARP1 PPP4R2 CDC7 MCM5 LIG3 MCM2 MCM4 GINS2 FEN1 MCM3 NUCKS1 MCM6 RPA3 MCM7 SAMHD1 RPA1 MMS19 UBE2N GINS4 PPP4C RPA2 RECQL YY1 NSMCE2 NBN EXD2 CDC45 ACTR2 RAD51 RAD50 MRE11 RIF1 CGAS NSMCE1 NIPBL SFPQ INO80 PALB2 WDR48 TP53BP1 SMC5 AP5Z1 BLM TERF2IP RMI1

Figure S3

***Figure S3 Loss of USP28 negatively affects the expression of DDR effector proteins in SCC***

A) Expression of DNA damage gene expression according to Kauffmann signature selection in human lung squamous cell carcinomas (SCC, n=486), adenocarcinomas (ADC, n=483) and normal non-transformed tissue (normal SCC=50, normal ADC=59). Generated with the open source tool www.gepia2.cancer-pku.cn. In box plots, the center line reflects the median, the upper and lower box limits indicates the first and third quartile. Whiskers extend 1.5x the IQR and outliers are marked as dots.

B) Volcano Plots with the expression of the gene signatures Shedden lung cancer poor survival A4 (left panel) and Shedden lung cancer good survival A6 (right panel) comparing lung ADC and SCC patients. Generated with the online tools www.r2.amc.nl.

C) Volcano plot proteome analysis of A431 cells treated with the DUB inhibitor AZ-1 or DMSO (Ctrl), shRNA targeting USP28 or Non-targeting control NTC (Ctrl). n=3

D) Spearman correlation of the proteome analysis between A431 cells treated with the DUB inhibitor AZ-1 or DMSO (Ctrl) and A431 sh-USP28 or Non-targeting control NTC (Ctrl). n=3

E) Heatmap proteome analysis according to the Shedden lung cancer poor survival signature of A431 cells treated with the DUB inhibitor AZ-1 or DMSO Control), shRNA targeting USP28 or Non-targeting control NTC (Control). n=3.

F) Whole heatmap proteome analysis of GO Recombinational Repair signature in control (ctrl) vs sh-USP28#1 and DMSO (ctrl) vs 15μM AZ1 A431 cells. Red = DNA damage sensors; Yellow = DNA damage sensor for NHEJ.

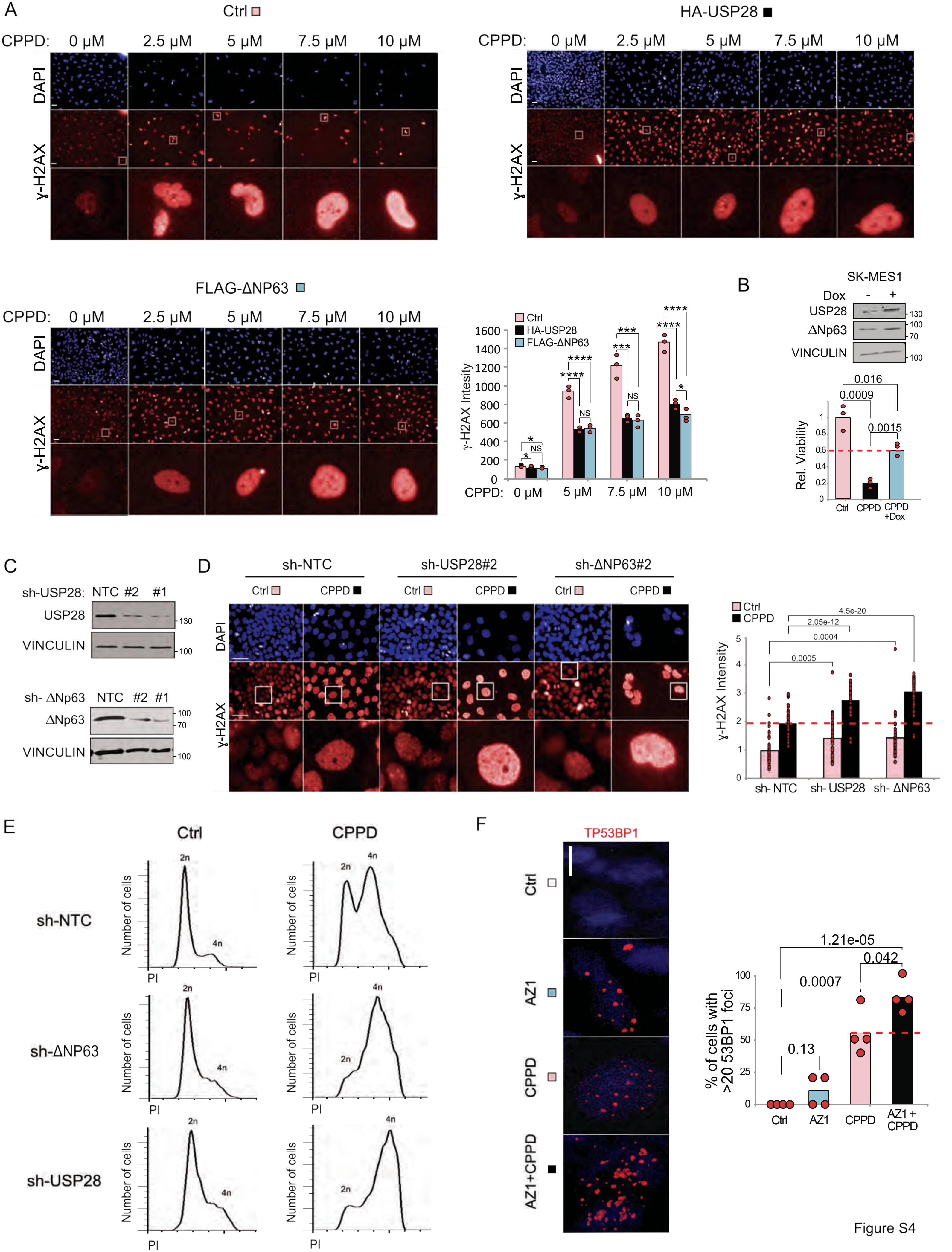
A
Ctrl
HA-USP28
FLAG-ΔNP63
CPPD: 0 μM 2.5 μM 5 μM 7.5 μM 10 μM
DAPI
γ-H2AX
Ctrl
HA-USP28
FLAG-ΔNP63
γ-H2AX Intesity
1600
1400
1200
1000
800
600
400
200
0
CPPD: 0 μM 5 μM 7.5 μM 10 μM
NS
B
SK-MES1
Dox - +
USP28
ΔNp63
VINCULIN
130
100
70
100
Rel. Viability
0.016
0.0009
0.0015
Ctrl
CPPD
CPPD +Dox
C
sh-USP28: NTC #2 #1
USP28
VINCULIN
sh- ΔNp63 NTC #2 #1
ΔNp63
VINCULIN
D
sh-NTC
sh-USP28#2
sh-ΔNP63#2
Ctrl
CPPD
DAPI
γ-H2AX
γ-H2AX Intensity
4.5e-20
2.05e-12
0.0004
0.0005
sh- NTC
sh- USP28
sh- ΔNP63
E
Ctrl
CPPD
sh-NTC
sh-ΔNP63
sh-USP28
Number of cells
PI
2n
4n
F
TP53BP1
Ctrl
AZ1
CPPD
AZ1+CPPD
% of cells with >20 53BP1 foci
1.21e-05
0.042
0.0007
0.13
100
75
50
25
0
AZ1 + CPPD

Figure S4

***Figure S4 USP28-ΔNp63 axis is required for DDR upon cisplatin treatment and chemoresistance in SCC***

A) Immunofluorescence staining against the DNA damage marker phospho-H2AX in BEAS-2B cells transiently transfected with either human USP28 or ΔNp63. Transfection of a GFP cDNA expressing plasmid served as control (-). Cells were exposed to indicated concentrations of CPPD for 48 hours. DAPI served as nuclear marker. n=3. Quantification of relative phospho-H2AX fluorescence intensity was measured quantifying 15 fields per well (n=3). Scale bar= 100μm White boxes were used in Fig 4B. P-values were calculated using two-tailed T-test statistical analysis.

B) Immunoblotting of USP28 and ΔNp63 in SK-MES1 cells lentivirally transduced with inducible mUSP28 upon exposure to either DMSO or 1μM DOX for 96 hours. VINCULIN serves as loading control. Relative viability was quantified using SK-MES1 cells lentivirally transduced with inducible mUSP28 upon exposure to DMSO, 15μM CPPD or 1μM DOX + 15μM CPPD for 96 hours. P-values were calculated using two-tailed T-test statistical analysis.

C) Immunoblot of USP28 and ΔNp63 in A431 cells lentivirally transduced with two constitutive shRNA targeting either USP28 or ΔNp63. VINCULIN serves as loading control. n=3.

D) Immunofluorescence staining against phospho-H2AX in lentivirally transduced A431 cells (shRNA-control, shRNA USP28#2 or ΔNp63#2) upon exposure to either DMF or 5μM CPPD for 48 hours. DAPI served as nuclear marker. n=3. Quantification of relative phospho-H2AX fluorescence intensity in A431 cells. Scale bar= 200 μm n=50 cells. Two tailed T-Test was used to calculate the p-value

E) FACS-based cell cycle analysis in lentivirally transduced A431 cells (shRNA-control, shRNA USP28#1 or ΔNp63#1) upon exposure to either DMF or 5μM CPPD for 48 hours. n=3.

F) Immunofluorescence staining against TP53BP1 in A431 cells upon exposure to DMF+DMSO (Ctrl), 5μM CPPD, 15μM AZ1 or 5μM CPPD + 15μM AZ1 for 48 hours. DAPI served as nuclear marker. Percentage of cells with more than 20 P53BP1 foci was calculated measuring 10 cells per field (n=5). Scale bar= 10μm. P-values were calculated using two-tailed T-test statistical analysis.

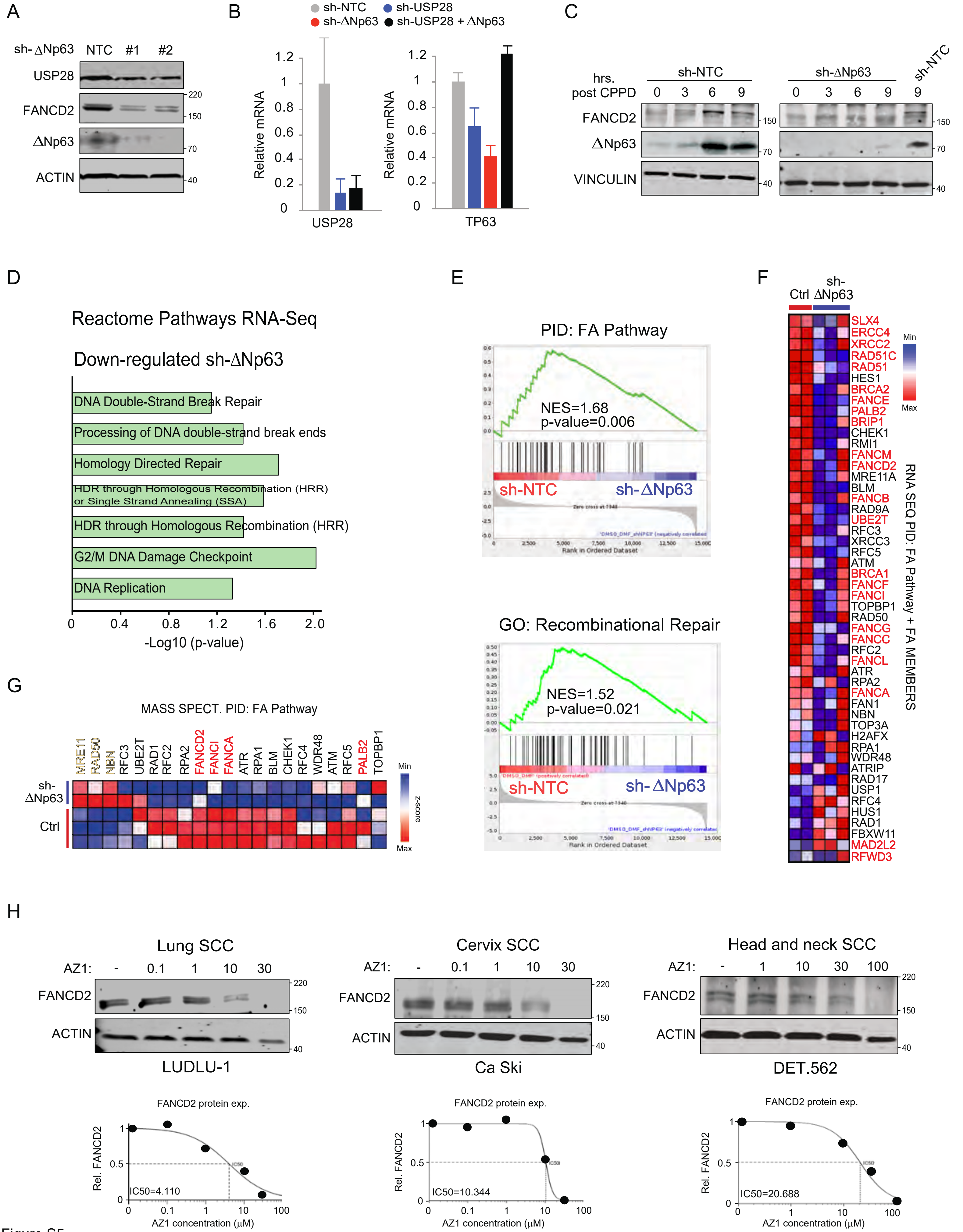

A
sh- ΔNp63 NTC #1 #2
USP28
FANCD2
ΔNp63
ACTIN
B
sh-NTC sh-USP28 sh-ΔNp63 sh-USP28 + ΔNp63
Relative mRNA
USP28
TP63
C
hrs. post CPPD
sh-NTC
sh-ΔNp63
FANCD2
ΔNp63
VINCULIN
D
Reactome Pathways RNA-Seq
Down-regulated sh-ΔNp63
DNA Double-Strand Break Repair
Processing of DNA double-strand break ends
Homology Directed Repair
HDR through Homologous Recombination (HRR) or Single Strand Annealing (SSA)
HDR through Homologous Recombination (HRR)
G2/M DNA Damage Checkpoint
DNA Replication
-Log10 (p-value)
E
PID: FA Pathway
NES=1.68
p-value=0.006
sh-NTC
sh-ΔNp63
GO: Recombinational Repair
NES=1.52
p-value=0.021
sh-NTC
sh-ΔNp63
F
Ctrl
sh-ΔNp63
Min
Max
SLX4 ERCC4 XRCC2 RAD51C RAD51 HES1 BRCA2 FANCE PALB2 BRIP1 CHEK1 RMI1 FANCM FANCD2 MRE11A BLM FANCB RAD9A UBE2T RFC3 XRCC3 RFC5 ATM BRCA1 FANCF FANCI TOPBP1 RAD50 FANCG FANCC RFC2 FANCL ATR RPA2 FANCA FAN1 NBN TOP3A H2AFX RPA1 WDR48 ATRIP RAD17 USP1 RFC4 HUS1 RAD1 FBXW11 MAD2L2 RFWD3
RNA SEQ PID: FA Pathway + FA MEMBERS
G
MASS SPECT. PID: FA Pathway
MRE11 RAD50 NBN RFC3 UBE2T RAD1 RFC2 RPA2 FANCD2 FANCI FANCA ATR RPA1 BLM CHEK1 RFC4 WDR48 ATM RFC5 PALB2 TOPBP1
sh-ΔNp63
Ctrl
Min
Max
z-score
H
Lung SCC
AZ1: - 0.1 1 10 30
FANCD2
ACTIN
LUDLU-1
FANCD2 protein exp.
Rel. FANCD2
IC50=4.110
AZ1 concentration (μM)
Cervix SCC
AZ1: - 0.1 1 10 30
FANCD2
ACTIN
Ca Ski
FANCD2 protein exp.
Rel. FANCD2
IC50=10.344
AZ1 concentration (μM)
Head and neck SCC
AZ1: - 1 10 30 100
FANCD2
ACTIN
DET.562
FANCD2 protein exp.
Rel. FANCD2
IC50=20.688
AZ1 concentration (μM)


Figure S5

***Figure S5 Deregulation of ΔNp63 impairs the Fanconi Anemia pathway in SCC***

A) Immunoblot of endogenous USP28, FANCD2 and ΔNP63 in A431 cells lentivirally transduced with constitutive shRNA targeting either control or two independent sequences targeting ΔNP63. ACTIN serves as loading control. n=3.

B) Quantitative RT-PCR of *USP28 and TP63* in A431 sh-NTC, sh-USP28#1 & #2, sh-ΔNp63#1 & #2 or sh-USP28#1 transfected with ΔNp63 cells normalised to ACTB. Quantitative graphic is represented as mean ± SD of three experiments (n= 3).

C) Immunoblot of FANCD2 in 1 hours treated CPPD chase experiment (5μM) of control (sh-NTC) and sh-ΔNP63#1 A431 cells. Numbers indicate hours post CPPD treatment. VINCULIN served as loading control. n=3.

D) Downregulated Reactome pathways related to DNA damage signalling upon depletion of ΔNP63 in A431 cells. Generated with the open source tool www.pantherdb.org

E) Gene set enrichment analysis (GSEA) of FA pathway signature genes and Recombinatorial Repair pathway signature genes in sh-ΔNP63#1 infected A431 cells. The gene set was analysed in sh-NTC (left) and sh-ΔNP63#1 depleted (right) A431 cells. (*N*)ES: (normalized) enrichment score.

F) RNA sequencing heatmap of FA pathway and pathway member genes in control or sh-ΔNP63#1 silenced A431 cells. Direct FA members are highlighted in red.

G) Heatmap of whole proteome analysis of FA pathway signature genes in control or sh-ΔNP63#1 silenced A431 cells. Direct FA members are highlighted in red.

H) Immunoblot of endogenous FANCD2 in the SCC cell lines LUDLU-1 (LSCC), Ca Ski (CESC) and Detroit 562 (HNSCC) cells treated for 24 h with either DMSO or indicated concentrations of AZ1. ACTIN served as loading control. FANCD2 half-maximal inhibitory protein abundance ($IC_{50}$) was calculated.

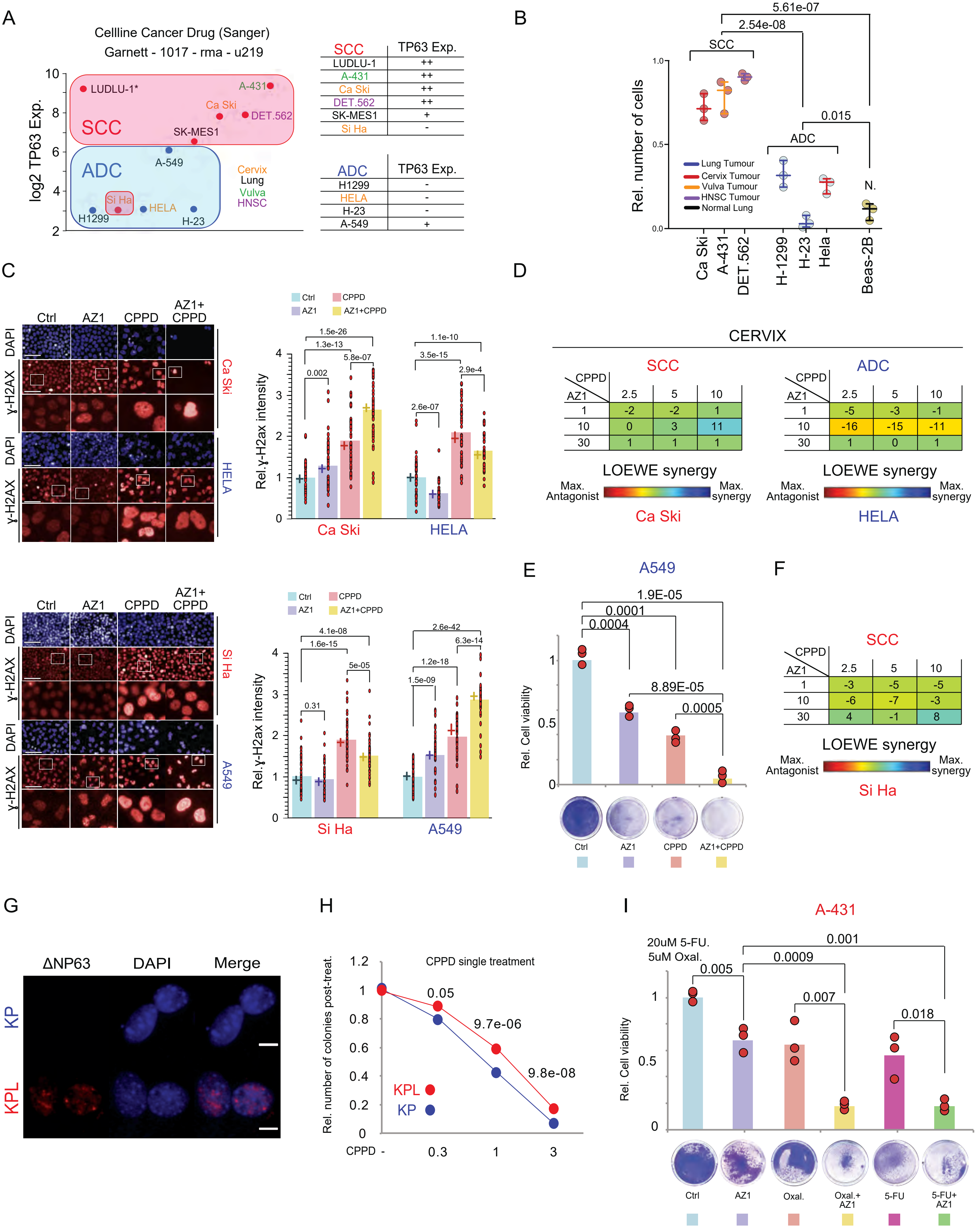


D — SCC (Ca Ski), LOEWE synergy

| AZ1 \ CPPD | 2.5 | 5 | 10 |
|---|---|---|---|
| 1 | -2 | -2 | 1 |
| 10 | 0 | 3 | 11 |
| 30 | 1 | 1 | 1 |

D — ADC (HELA), LOEWE synergy

| AZ1 \ CPPD | 2.5 | 5 | 10 |
|---|---|---|---|
| 1 | -5 | -3 | -1 |
| 10 | -16 | -15 | -11 |
| 30 | 1 | 0 | 1 |

F — SCC (Si Ha), LOEWE synergy

| AZ1 \ CPPD | 2.5 | 5 | 10 |
|---|---|---|---|
| 1 | -3 | -5 | -5 |
| 10 | -6 | -7 | -3 |
| 30 | 4 | -1 | 8 |

Figure S6

***Figure S6 Pharmacologic inhibition of USP28 re-sensitizes SCC cells to chemotherapy***

A) Classification of human cancer cell lines of various origins relative to ∆NP63 expression status. Red box= SCC; Blue box= ADC. The expression of the different cell lines except LUDLU1 (*) were obtained from the dataset: Cellline Cancer Drug (Sanger) Garnett - 1017 - rma - u219. * = LUDLU1 gene expression was obtained from Cell line CCLE Cancer Cell Line Encyclopedia dataset. Data generated with the open source tool www.r2.amc.nl

B) Relative number of cells in A431, Ca Ski, Detroit 562, H-1299, H23, HeLa and Beas-2B cell lines after exposure to 2.5µM CPPD for 96 hours. P-values were calculated using two-tailed T-test statistical analysis.

C) Immunofluorescence staining of phospho-H2AX in Ca Ski, HeLa, Si Ha and A549 cell lines treated with DMSO+DMF (Ctrl), 15µM AZ1, 5µM CPPD or 15µM AZ1+5µM CPPD for 48 hours. DAPI served as nuclear marker. Relative quantification of the phospho-H2AX staining intensity was measured for the different treatment exposures. n= 50 cells. Scale bar= 200µm. P-values were calculated using two-tailed T-test statistical analysis. Red= SCC cell line; Blue= ADC cell line.

D) LOEWE synergism score of CPPD and AZ1 in Ca Ski and HeLa cell lines. Cells were exposed to the Indicated concentrations (µM) for 48 hours. DAPI was used to assess total cell numbers. Red= SCC cell line; Blue= ADC cell line.

E) Relative cell viability in A549 cells upon treatment with DMSO+DMF (Ctrl), 15µM AZ1, 5µM CPPD or 15µM AZ1+5µM CDDP for 48 hours. n=3. P-values were calculated using two-tailed T-test statistical analysis.

F) LOEWE synergism score of CPPD and AZ1 in Si Ha cells. Cells were exposed to the Indicated concentrations (µM) for 48 hours. DAPI was used to assess total cell numbers. Red= SCC cell line; Blue= ADC cell line.

G) Immunofluorescence staining of ∆NP63 in KP and KPL mouse cell lines. DAPI served as nuclear marker. Red= SCC cell line; Blue= ADC cell line. Scale bar= 10µm.

HI) Relative number of colonies upon exposure to indicated concentration of CPPD in KP and KPL cell lines. n=11. P-values were calculated using two-tailed T-test statistical analysis.

I) Relative cell viability in A431 cells upon treatment with DMSO+DMF (Ctrl), 15µM AZ1, 5µM Oxaliplatin, 15µM 5-FU, 15µM AZ1+5µM Oxaliplatin or 15µM AZ1+15µM 5-FU for 48 hours. n=3. P-values were calculated using two-tailed T-test statistical analysis.

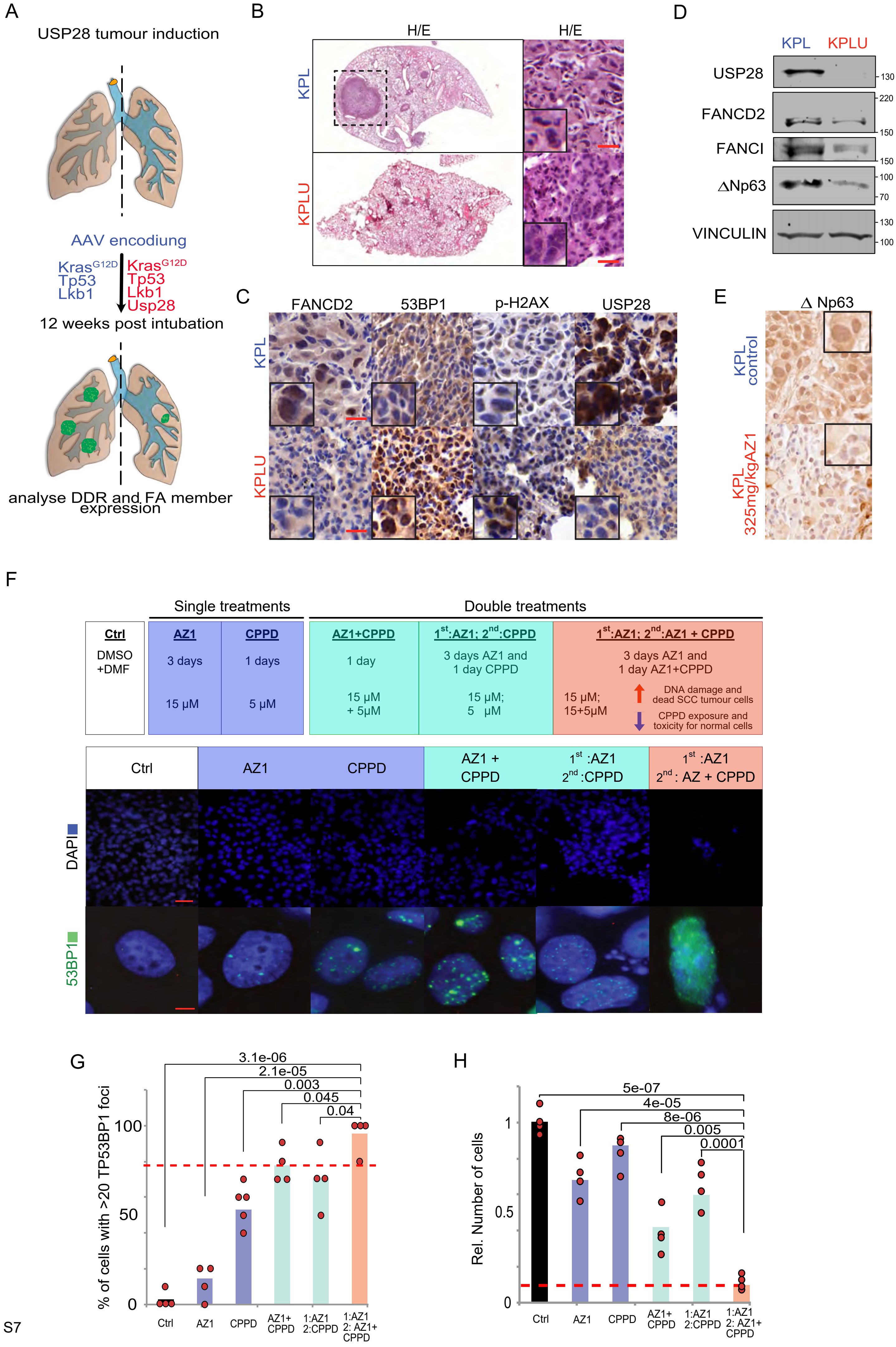
A
USP28 tumour induction
AAV encodiung
KrasG12D
Tp53
Lkb1
KrasG12D
Tp53
Lkb1
Usp28
12 weeks post intubation
analyse DDR and FA member expression
B
H/E
H/E
KPL
KPLU
D
KPL
KPLU
USP28
FANCD2
FANCI
ΔNp63
VINCULIN
130
220
150
150
100
70
130
100
C
FANCD2
53BP1
p-H2AX
USP28
KPL
KPLU
E
Δ Np63
KPL control
KPL 325mg/kgAZ1
F
Single treatments
Double treatments
Ctrl
DMSO +DMF
AZ1
3 days
15 µM
CPPD
1 days
5 µM
AZ1+CPPD
1 day
15 µM + 5µM
1st:AZ1; 2nd:CPPD
3 days AZ1 and 1 day CPPD
15 µM; 5 µM
1st:AZ1; 2nd:AZ1 + CPPD
3 days AZ1 and 1 day AZ1+CPPD
15 µM; 15+5µM
DNA damage and dead SCC tumour cells
CPPD exposure and toxicity for normal cells
Ctrl
AZ1
CPPD
AZ1 + CPPD
1st :AZ1 2nd :CPPD
1st :AZ1 2nd : AZ + CPPD
DAPI
53BP1
G
% of cells with >20 TP53BP1 foci
3.1e-06
2.1e-05
0.003
0.045
0.04
100
50
0
Ctrl
AZ1
CPPD
AZ1+ CPPD
1:AZ1 2:CPPD
1:AZ1 2: AZ1+ CPPD
H
Rel. Number of cells
5e-07
4e-05
8e-06
0.005
0.0001
1
0.5
0
Ctrl
AZ1
CPPD
AZ1+ CPPD
1:AZ1 2:CPPD
1:AZ1 2: AZ1+ CPPD

Figure S7

***Figure S7 Inhibition of USP28 activity deregulates FA-DDR signalling in vivo and sensitizes tumours to CPPD treatment in ex vivo organotypic lung SCC tumor slice cultures by de-activating FA***

A) Schematic diagram of CRISPR/Cas9-mediated tumour modelling and targeting of *p53*$^{\Delta}$*; Lkb1*$^{\Delta}$*; KRas*$^{G12D}$(KPL) or *Usp28*$^{\Delta}$*; p53*$^{\Delta}$*; Lkb1*$^{\Delta}$*:KRas*$^{G12D}$(KPLU) in *Rosa26Sor-CAGG-Cas9-IRES-GFP* mice.

B) Representative images of sections from KPL and KPLU mice stained with haematoxilin and eosin. Inlay shows higher magnification.

C) Immunohistochemistry of FANCD2, TP53BP1, phospho-H2AX and USP28 in primary KPL or KPLU tumors. Scale bar = 50µm

D) Immunoblot of endogenous USP28, FANCD2, FANCI and ∆Np63 from KPL or KPLU tumours. VINCULIN served as loading control. n = 3.

E) Immunohistochemistry of ∆Np63 in primary KPL tumors from vehicle or AZ1 treated mice. Scale bar = 50µm

F) Upper panel= Schematic representation of treatment strategies combining AZ1 (15 µM) with CPPD (5µM) in A431 cells. Lower panel= Immunofluorescence staining of the DNA damage marker P53BP1 in A431 cells treated with compounds and times according to upper panel, respectively. DAPI serves as nuclear staining control. n=3. Upper scale bar = 100 µm; Lower scale bar= 10 µm.

G) Quantification of % cell with >20 TP53BP1 nuclear foci in A431 cells treated according to F). TP53BP1 foci were calculated measuring 10 cells per field (n=5). P-values were calculated using two-tailed T-test statistical analysis.

H) Relative surviving number of cells in A431 cells treated according to F). n=3. P-values were calculated using two-tailed T-test statistical analysis.
